# Robust Population Single Neuronal Calcium Signal Extraction Using SCOUT Allows for Longitudinal Analysis of Behavior-associated Neural Ensemble Dynamics

**DOI:** 10.1101/2020.08.26.268151

**Authors:** Kevin G. Johnston, Steven F. Grieco, Zhaoxia Yu, Suoqin Jin, Tong Shen, Rachel Crary, John F. Guzowski, Todd C. Holmes, Qing Nie, Xiangmin Xu

**Author notes:** **Correspondence should be addressed to** Dr. Qing Nie, Department of Mathematics, University of California, Irvine, CA, and Dr. Xiangmin Xu, PhD, Department of Anatomy and Neurobiology, School of Medicine, University of California, Irvine, CA 92697.

## Abstract

*In vivo* calcium imaging enables simultaneous recording of large neuronal ensembles while engaged in operations such as learning and memory. However, such *in vivo* optical recordings are typically subject to motion artifact and background contamination from neurons and blood vessels. Further, population cell tracking across multiple recordings is complicated by non-rigid transformation induced by cell movements and field shifts. We introduce the novel method SCOUT for Single-Cell SpatiOtemporal LongitUdinal Tracking, consisting of two crucial parts: (1) imposition of spatial constraints on neuronal footprints extracted from individual optical recordings to improve ROI selection and eliminate false discoveries, and (2) application of a predictor-corrector, using spatiotemporal correlation of extracted neurons across sessions, for population cell tracking across multiple sessions. SCOUT empirically outperforms current methods for cell extraction and tracking in long-term multi-session imaging experiments across multiple brain regions. Application of this method allows for robust longitudinal analysis of contextual discrimination associated neural ensemble dynamics in the hippocampus up to 60 days.

## Introduction

Extracting longitudinal activity of large-scale neuronal ensembles is a fundamental first step to the analysis of neural circuit responses. Ca++ imaging of population neurons allows the recording of larger neural ensembles than can be recorded electrically. *In vivo* calcium imaging using microendoscopic lenses enables imaging of previously inaccessible large ensembles of neuronal populations at single-cell level in freely moving mice as they experience environmental stimuli, and perform neural transformations that underlie behavioral responses over both short and long timescales ([Flusberg et al., 2008], [Ghosh et al., 2011], [Ziv and Ghosh, 2015]). Microendoscopic *in vivo* brain imaging via head-mounted fluorescent miniature microscopes (“miniscopes”) have been used to study a diverse set of neural circuits in the hippocampus ([Cai et al., 2016], [Ziv et al., 2013], [Jimenez et al., 2016], [Rubin et al., 2015], [Sun et al., 2019]), entorhinal cortex ([Kitamura et al., 2015], [Sun et al., 2015]), striatum ([Barbera et al., 2016], [Klaus et al., 2017]) and amygdala [Yu et al, 2017], among other regions. This powerful technique promises to open the “black box” of critical neural transformations in higher level brain areas that occur between sensory inputs and motor outputs.

New advances in 1-photon (1p) miniscope imaging data analysis increase our capability to study neural ensembles, but require the extraction of neural activity from recordings, which consists of the isolation of temporal signals (fluorescence traces), and spatial footprints (pixel intensity values) corresponding to the individual neurons in the recording. Isolating such signals from the recording is difficult due to spatial overlaps between neurons, and complicated background effects that interfere with the neuronal temporal signal.

Several methods have been developed to extract neural activity recorded as optical imaging signals ([Apthorpe et al., 2016], [Mukamel et al., 2009], [Pnevmatikakis et al., 2016], [Zhou et al., 2018], [Giovannucci et al., 2019], [Pnevmatikakis, 2019]). Briefly, such methods can be divided into those which estimate regions of interest (ROIs) for each neuron, followed by extraction of the temporal signal, and methods which iteratively update both temporal and spatial components of the individual neurons. Recently, nonnegative matrix factorization methods such as CNMF and its 1-photon variant CNMF-E ([Pnevmatikakis et al., 2016; Zhou et al., 2018]), members of the latter category, have been extensively used for neural signal extraction from optical recordings ([Trevathan et al., 2018], [Gonzalez et al., 2019]) (see **Materials and Methods** and [Pnevmatikakis, 2019] for additional details).

One major issue for current extraction methods is the prevalence of false discoveries, which consist of extracted footprints and temporal traces that do not correspond to ground truth neurons in the recording (Supplementary Fig. 1). These false discoveries can be caused by background noise, inaccurate initialization of neuron footprints, or errors in the estimation of footprints and corresponding temporal traces. For *in vivo* recordings, depending on recording quality and initialization parameter, false discovery rates may reach 45% of detected neurons.

Some approaches have been suggested to address this issue, such as manual false discovery identification [Zhou et al. 2018], and convolutional neural network classification [Giovannucci et al., 2019], performed at intermediate stages of the extraction. However, for experiments involving large numbers of recordings, manual identification of false discoveries becomes untenable. Convolutional neural networks have been used to detect false discoveries, but they lack interpretability, and struggle to generalize to neurons with different footprint profiles than those in the training set (see **Materials and Methods** for additional discussion).

False discoveries in individual recordings introduce difficulties in studying the evolution of neural dynamics over time, as cells must be identified across multiple recording sessions, and false discoveries can interfere with the identification of cells across recordings. Assuming accurate individual recording extractions, attempts to extract the activity of neurons over long term experiments have taken one of three forms (Fig. 1A). (1) Initial concatenation of registered recordings followed by extraction of fluorescence traces and spatial footprints from the concatenated recording [Sun et al, 2019]. (2) Spatial patch methods [Zhou, et al. 2018] in which registered recordings are concatenated, and the spatial dimension is split into overlapping patches. Extraction is performed on each patch separately, and neurons are merged across the patches, giving extracted footprints and traces for the entire recording. (3) Temporal batch methods such as cellReg [Sheintuch et al., 2017], in which footprint and fluorescence traces are extracted and cells are tracked across multiple recordings using spatial similarity.

**Figure 1.**
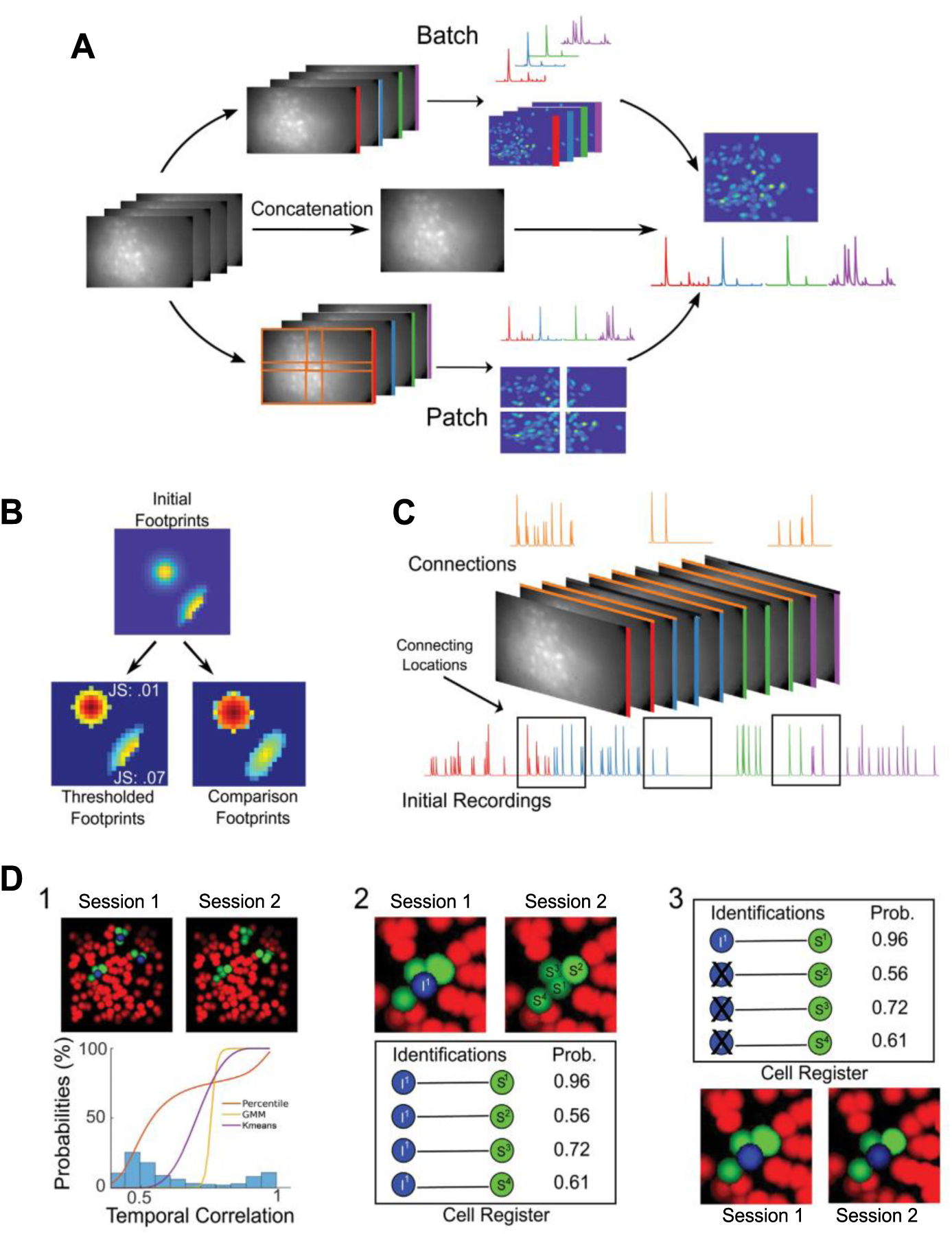
Schematic overview of SCOUT. **A:** Workflow for extracting fluorescence traces and spatial footprints from multiple sessions include spatial patch methods (bottom track), concatenation (center track) and temporal batch methods (top track). Our method, SCOUT, is a modified temporal batch method. **B:** Application of spatial constraints allows us to detect significant deviation from proposed footprint shapes. An individual neuron (top) is thresholded (bottom left) based on pixel intensity. The individual components are compared with baselines from a parameterized family of distributions (bottom right). JS divergence measures information loss in estimating extracted shape (bottom left). with proposed shapes (bottom right), showing increased information loss in estimating the footprint of the bottom right neuron. The bottom right footprint was created by cropping a true footprint, so the increase in JS divergence indicates correctly that the neuron should be removed from consideration. **C**: Connecting recordings (orange; middle) are created from selecting a subset of frames from subsequent recordings (red, blue, green, purple; middle). Fluorescence traces from connecting recordings (top) are used to identify fluorescence traces throughout the whole recording that correspond to a single neuron (bottom). Boxed regions of the full fluorescence trace (lower) indicate areas of comparison with connecting recordings. **D**: Illustration of cell tracking portion of SCOUT: (1) From the spatial footprints (colored ovals) of extracted neurons from an initial recording (top left), targeted neurons for tracking across sessions is selected (blue), and their surrounding neighbors determined (green). In the secondary recording (top right), we identify possible choices for tracking between sessions (green) using a maximal distance between centroids in the proposed choices in the secondary recording, and that of the neurons targeted for tracking. Weighted similarities (histogram) between all possible neighboring pairs of neurons between sessions are calculated. The x-axis indicates the weighted probabilities of possible neighbors, and the y-axis indicates normalized counts of probabilities between possible neighbors. The colored curves indicate possible functions for assigning tracking probabilities to associated pairs of neurons. Methods of probability assignment include percentile, soft K-means, and Gaussian Mixture Models. (2) top: We zoom in on one initial neuron (I^1^), and its possible identifications in the secondary recording. bottom: Possible neuron chains are constructed, and chain probabilities calculated for each. (3): top: Duplicate neurons in the first entry of each chain are eliminated based on the chain probability. bottom: The remaining neuron chain indicates that S^1^ is the most probable identification for I^1^ in the secondary recording.

Concatenation and patch methods can be resource intensive in terms of both computational power and time requirements. In terms of scalability, temporal batch methods (cell tracking methods) may provide the best option for long-term neural ensemble analysis, but existing methods do not track many available neurons due to exclusive use of spatial metrics for determining inter-recording neuron similarity.

To address these issues, we present **SCOUT** (Single-Cell SpatiOtemporal LongitUdinal Tracking), an end-to-end modular system for accurate extraction and tracking of individual neurons in long-term experiments. We demonstrate the effectiveness of SCOUT on both simulated and experimental data from multiple brain regions including the hippocampus, the visual cortex, and the prefrontal cortex within individual sessions and across multiple sessions. Moreover, we track single cell populations through long-term contextual discrimination experiments in mouse hippocampal CA1 up to 60 days with *in vivo* GCaMP6-based calcium imaging in freely moving animals. This enables us to analyze the evolution of context dependent neural ensembles through learning, extinction, and relearning phases of a contextual discrimination experiment with better performance than previously available methods.

## Results

### SCOUT overview

Long-term experiments require both the detection of neurons from individual recordings, and cell tracking, in which neurons are tracked through each recording. SCOUT consists of two major modules: <u>individual recording extraction</u>, and <u>cell tracking to identify the same neurons</u> <u>across multiple sessions</u>.

An initial preprocessing step starts with motion correction preprocessing of recordings using NoRMCorre [Pnevmatikakis and Giovannucci, 2017], followed by registration of recording sessions. SCOUT provides several methods for template extraction and image registration, as well as an interface for manual interventions, if initial automatic results are unsatisfactory.

### SCOUT: individual recording extraction

The individual recording extraction module of SCOUT consists of a neural signal extraction algorithm (such as CNMF-E), to which we have added a binary classifier for identifying false discoveries. Motivated by our previous success of using approximate matched filters for automatically detecting and extracting neural synaptic inputs [Shi et al., 2010], we introduce a spatial template filter, which eliminates false discoveries and improves fluorescence trace quality via imposition of spatial constraints based on a proposed baseline (Fig. 1B; Supplementary Fig. 1). We assume (normalized) spatial footprints for detected neurons are members of a family of two-dimensional probability distributions, and compute a metric indicating the similarity between proposed spatial footprints, and the closest neighbor in the family of distributions. Neurons with large divergence from the family of distributions are deleted, and the proposed footprints of retained neurons may be updated via multiplication with the binary mask of the compared distribution (**Algorithm 1, Supplementary Video 1)**. Similarity between distributions is calculated using Jensen-Shannon divergence.

In practice, we suggest running the filter at each iteration after spatial components are updated, but before the temporal components are updated, to maximize signal extraction accuracy (Supplementary Fig. 1G). The spatial filter may also be used as a post-processing step after neuron extraction, allowing for easy inclusion in any pipeline.

Regarding appropriate families of distributions for 1-photon data, we have considered a Gaussian model, in which the centroid and covariance is extracted from the proposed footprint, to construct a similar bivariate normal distribution, and an elliptical model, in which the centroid, major axis length, minor axis length, and orientation are used to construct an elliptical footprint, with pixel intensity values modeled using the rate of intensity decrease from centroid to boundary of the proposed neuron. Experimentally, the elliptic model seems to provide a better baseline for false discovery identification (see **Materials and Methods**, Supplementary Fig. 1F). Possible future extensions include a ring model for 2-photon data.

### SCOUT: cell tracking

The second module of SCOUT deals with accurate single cell tracking across multiple sessions (see **Algorithm 2, Supplementary Video 2**). SCOUT uses the combination of spatial footprint location with extracted fluorescence traces to track neurons, via the construction of connecting segments between recordings (Fig. 1C). This allows for both spatial and temporal criteria to be used for neuron identification.

This module consists of three steps: (1) similarity probabilities are calculated between each session, (2) neurons are tracked through the full set of recordings, (3) a correction phase in which duplicate neurons are removed. The first two steps constitute a predictor, while the last step constitutes a corrector.

#### Algorithm 1 SCOUT Spatial Template Filter Overview

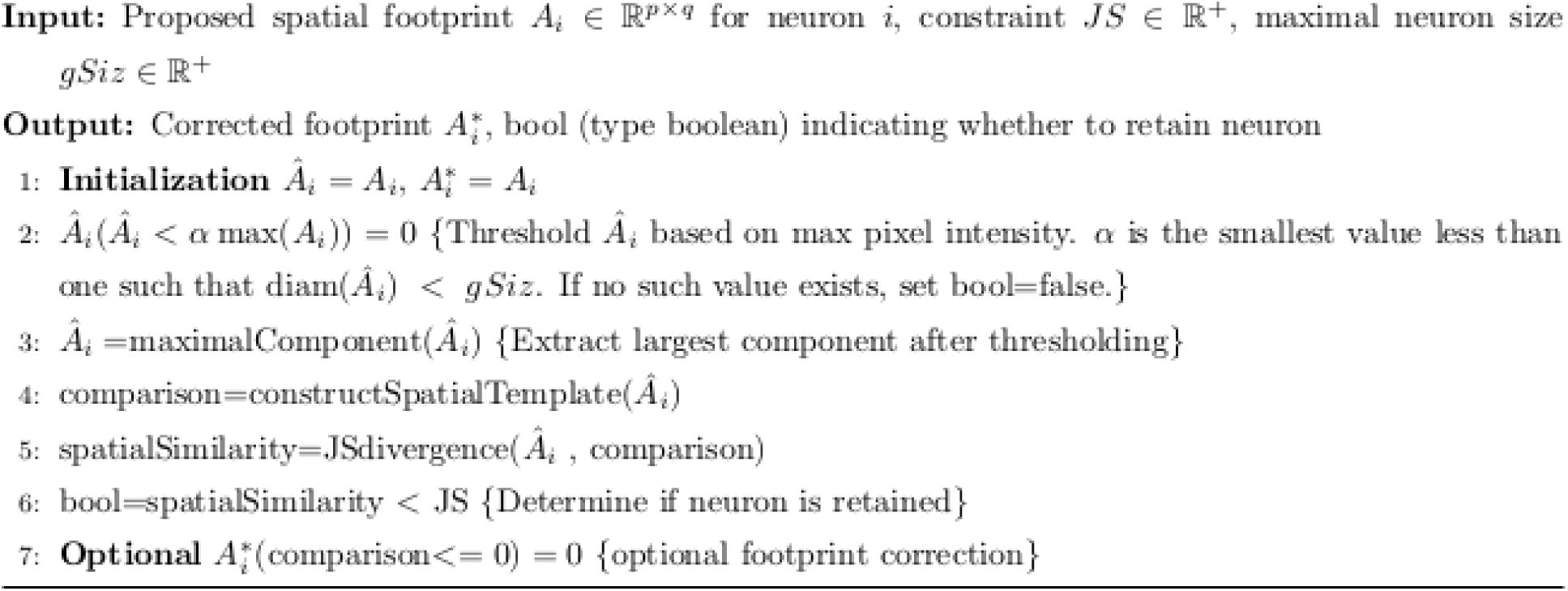

#### Algorithm 2 SCOUT Cell Tracking Overview

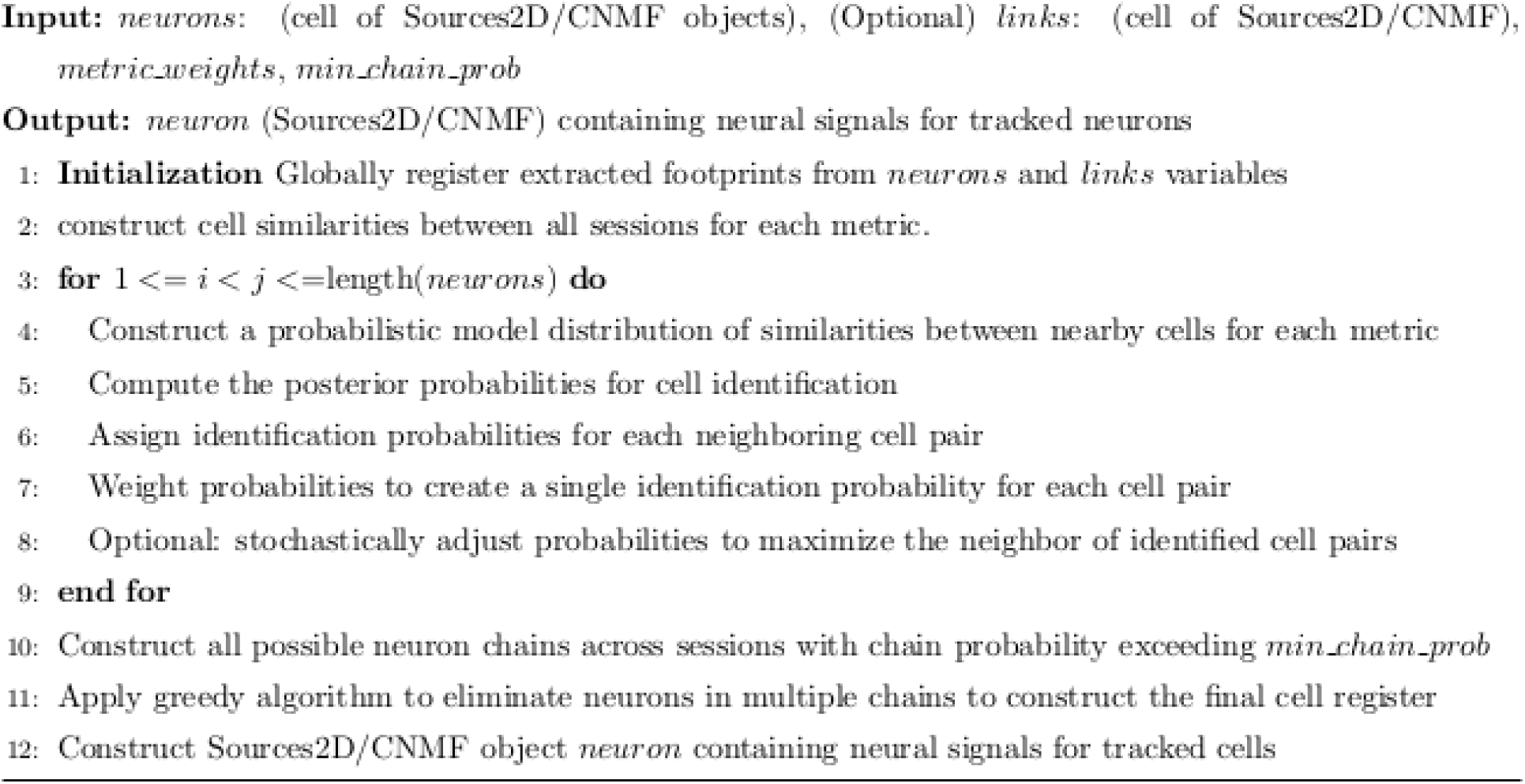

For each pair of consecutive recordings, a connecting recording is constructed, and spatiotemporal similarity scores (centroid distance, JS divergence, temporal correlation on overlap, signal-to-noise ratio, etc.) are calculated for neurons across the recording. Next, similarity scores are weighted to produce a single score for each pair of neurons from consecutive sessions (producing a weighted ensemble of predictors), and neuron identification probabilities between sessions are estimated using probabilistic models (Fig. 1D panel 1, Supplementary Fig. 1B, **Materials and Methods**). Methods of probability assignment (percentile, soft K-means, Gaussian Mixture Model) have various properties. Percentile and soft K-means assignment have wide parameter ranges on which they give non-negligible probabilities, while GMM typically gives a sharper boundary. GMM typically produces few false discoveries but can restrict the number of detected neurons. We suggest using soft K-means in many instances as identification errors are corrected via the weighted ensemble of predictors, and the correction phase ([Everitt, 2014], [Dunn, 1973]).

Neurons are tracked throughout the full set of recordings using the estimated probabilities, creating “chains” of neurons, where each chain contains at most one neuron extracted from each recording. Chain probabilities are calculated as the number of identified neurons in each chain (i.e. that exceed a certain similarity probability threshold), divided by the total number of possible pairs in the chain.

Chain probability estimation is followed by a correction phase in which a neuron extracted from a given recording is constrained to appear in at most one chain, by removing duplicate neurons from chains based on their identification probabilities (see Fig. 1D panel 3, Supplementary Fig. 2C, **Materials and Methods**). The finalized neuron chains are assembled into a cell register, a matrix in which each row contains the associated session identification numbers for a neuron chain. Because of the correction phase, columns of the cell register do not contain duplicate ids. The cell register is used to construct concatenated temporal signals corresponding to each tracked neuron; the cell register and the associated concatenated neural signals are the final output.

SCOUT differs from alternative cell tracking software in several ways. First, SCOUT incorporates temporal correlation similarity across an overlapping portion of each consecutive session. Second, SCOUT can incorporate an arbitrary number of similarity metrics into its calculations for neuron identification. Finally, exclusivity, a requirement that individual neurons appear in exactly one chain, is guaranteed by the correction phase of SCOUT.

### Comparisons of SCOUT with CNMF-E on simulated single session recordings

We tested the individual recording extraction portion of SCOUT on two different datasets: first, a set of 14 simulated recordings with 2000-10000 frames, consisting of between 50 and 200 neurons with Gaussian spatial footprints, plus simulated low-level noise and a simulated blood vessel; second, a dataset consisting of 40 recordings with 2000-8000 frames, in which initial Gaussian footprints were simulated, then altered via randomly generated non-rigid transformations, to simulate possible spatial distortions. In the second dataset, noise levels were higher compared with the true signal and blood vessel signal (see **Materials and Methods** for full simulation details). Hereafter, the first simulated dataset will be referred to as the Gaussian dataset, while the second simulated dataset will be referred to as the Non-rigid dataset. (Fig. 2, **Materials and Methods**).

**Figure 2.**
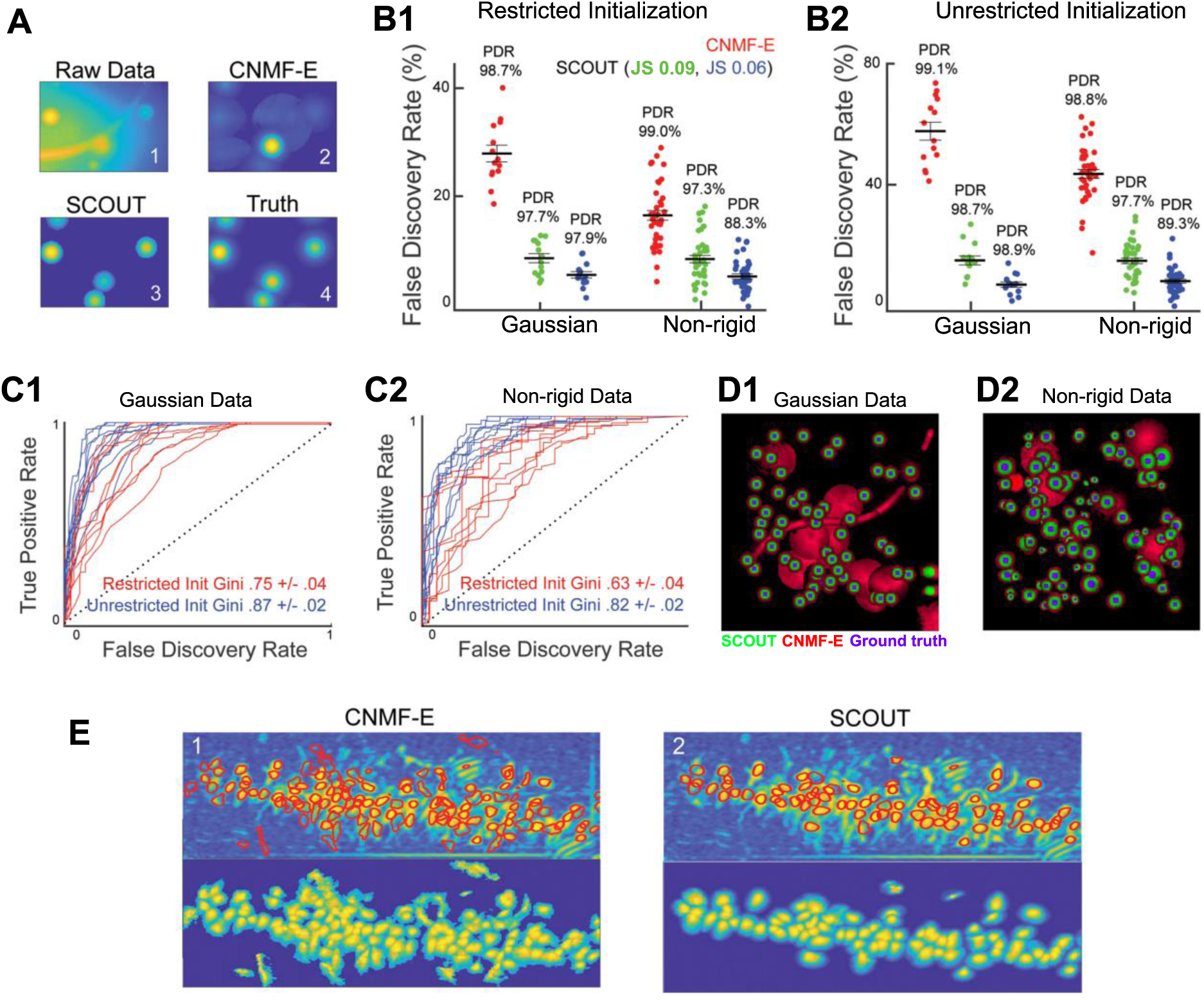
A cell-shape based spatial constraint improves extraction performance by controlling false discovery detection and improving neural extraction quality. **A**: Spatial footprint extraction quality shows improvement using SCOUT over CNMF-E on a simulated video recording (6000 frames). (1) maximum projection of a 70 pixel × 100 pixel section of a recording, (2) spatial footprint extraction results of CNMF-E applied to recording, showing multiple false discoveries (3) spatial footprint extraction results of SCOUT applied to recording, retaining all ground truth neurons with no false discoveries, (4) ground truth spatial footprints. **B**: False discovery rates across the Gaussian and Non-rigid datasets, displaying CNMF-E and SCOUT extraction results. (1) The results for extractions with restricted initialization, (2) results for unrestricted initialization. SCOUT has significantly fewer false discoveries than CNMF-E in both contexts, while maintaining comparable percent detected rates. **C:** Applying the SCOUT spatial filter after neuron initialization allows us to test its efficacy as a binary classifier, labeling neurons as true or false discoveries. Varying the spatial threshold, we plot the ROC curves and calculate the GINI index (defined as 2 × AUC −1, where AUC denotes the area under the curve for a given ROC curve) as a qualitative measurement of classifier efficiency. (1) The results on a set of 10 recordings from the Gaussian dataset, (2) a set of 10 recordings from the Non-rigid dataset. The high GINI indices indicate a robust classifier. **D**: Extracted spatial footprints from sample recordings in the Gaussian (1) and Non-rigid (2) datasets demonstrate the difference in false discoveries using SCOUT over CNMF-E. Neuron footprints are normalized to have the same maximal intensity for comparison purposes. Neurons are colored by which methods detected each extracted neuron, with SCOUT: green, CNMF-E: red, Ground Truth: Blue. CNMF-E detects background sources and blood vessels that do not correspond to ground truth neurons. **E**: Examples of extracted footprints from recordings of CA1 layer of the mouse hippocampus, conducted using CNMF-E (1), and SCOUT (2), (7178 frames sampled at 15 Hz). (1) top: The correlation image of the recording, with circled neurons corresponding to those detected by CNMF-E. bottom: The extracted spatial footprints detected by CNMF-E, normalized to have the same maximum pixel intensity. (2) top: The correlation image of the recording, with circled neuron corresponding to those detected by SCOUT. bottom: The extracted spatial footprints detected by SCOUT. CNMF-E produced an increased false discovery rate and gives an unclear segmentation of neurons from the recording. Using the three criteria discussed in the main body, 28% of extracted neurons were labeled as false discoveries in the CNMF-E extraction, while only 9% were labeled as false discoveries in the SCOUT extraction, demonstrating improved performance. Spatial footprints extracted via SCOUT were smoothed during the final spatial template application.

We examined two extraction conditions for each dataset, one with restricted initializations, in which the threshold for neuron initialization was set sufficiently high to exclude most spurious initialization points (which can exist either due to random fluctuations in background noise, or incomplete subtraction of discovered neurons during the initialization procedure), and the other in which thresholds for neuron initialization were low. Neuron initialization is primarily based on local correlation and signal intensity (see **Materials and Methods, Supplementary Fig. 1B-D** for full details). Extractions with unrestricted initializations had many false discoveries, which allows us to demonstrate the robustness of SCOUT. We term these extractions as restricted and unrestricted through the remainder of this section.

We performed 6 extractions on each dataset (Gaussian, Non-rigid), for each initialization condition (restricted, unrestricted), 5 extractions in which we varied JS divergence thresholds for the spatial constraints (JS constraint values [0.03, 0.06, 0.09, 0.12, 0.15]), and a CNMF-E extraction. We found little significant increase in the number of detected neurons for thresholds exceeding 0.09, and a sharp drop off in detected neurons for thresholds lower than 0.06, so we report statistics from the extractions with these two parameters (see Fig. 2A-D, Table 1). Larger thresholds may be required on *in vivo* data, particularly in cell tracking applications, as false positives have a smaller effect on the result.

**Table 1:** Average PDR and FDR reported with restricted and unrestricted extraction methods applied in Gaussian versus Non-Rigid datasets. Highlighted in bold is the outcome that maximizes PDR/(1+FDR). Reported as mean ± standard error.

| Method | Restricted |  |  |  | Unrestricted |  |  |  |
| --- | --- | --- | --- | --- | --- | --- | --- | --- |
|  | Gaussian |  | Non-Rigid |  | Gaussian |  | Non-Rigid |  |
|  | PDR (%) | FDR (%) | PDR (%) | FDR (%) | PDR (%) | FDR (%) | PDR (%) | FDR (%) |
| CNMF-E | 98.7 ± 0.3 | 27.9 ± 1.5 | 99.0 ± 0.2 | 16.6 ± 0.9 | 99.1 ± 0.3 | 58.2 ± 2.9 | 98.8 ± 0.3 | 44.4 ± 1.4 |
| SCOUT: 0.06 | **97.9 ± 0.5** | **5.8 ± 0.6** | 88.3 ± 0.7 | 5.5 ± 0.5 | **98.9 ± 0.3** | **8.6 ± 0.8** | 89.3 ± 0.7 | 9.7 ± 0.3 |
| SCOUT: 0.09 | 97.7 ± 0.5 | 8.8 ± 0.8 | **97.3 ± 0.3** | **9.7 ± 0.7** | 98.7 ± 0.3 | 16.4 ± 1.4 | **97.7 ± 0.4** | **16.3 ± 0.9** |

Pearson correlation was calculated between extracted fluorescence traces and ground truth neurons. An extracted neuron was labeled as a false discovery if the maximum correlation value between its fluorescence trace and any ground truth neuron was smaller than 0.8. A ground truth neuron was counted as detected if the maximum correlation between its trace and that of an extracted neuron was at least 0.8. Statistical results were similar for higher thresholds.

For each dataset and initialization condition, we computed the false discovery rate (FDR), defined as the percentage of false discoveries out of the detected neurons, and percent detected rate (PDR), defined as the percentage of ground truth neurons detected in a simulated recording for the extractions given by CNMF-E, SCOUT (JS: 0.06), and SCOUT (JS: 0.09). Statistics were calculated based on PDR and FDR calculated on each recording in each dataset. Results are reported in the form mean ±standard error. Statistical tests were two sided, and Welch’s paired t-test and one-way ANOVA were used where stated.

Applying one-way ANOVA to the results from each dataset and initialization condition separately, we identified significant difference of average FDR using CNMF-E, and SCOUT with constraints 0.06 and 0.09 (*p* < 7.4 × 10^-18^, taken over all datasets and initialization conditions). Pairwise comparisons between CNMF-E and SCOUT showed SCOUT detected fewer false discoveries on average at both constraint levels across all datasets and conditions (t-test *p <* 1.3 × 10^-7^). Average FDR exceeded 40% on the unrestricted extractions, and 16% on restricted extractions, across datasets. SCOUT reduced the number of false discoveries by at least half, and up to 85%, while retaining the percent detected rate within 1-2 percentage points of CNMF-E. Total reported average PDR generally exceeded 97%, depending on the dataset, initialization, and extraction method. As expected, more false negatives were assigned by SCOUT in the Non-rigid dataset, requiring a higher JS threshold (0.09) to retain at least 97% of true neurons (see Fig 2B, Table 1).

To further investigate the efficacy of SCOUT as a classifier, we considered 10 recordings randomly taken from each dataset and condition, and constructed receiver operating characteristic curves, by applying the spatial constraint after neuron initialization (Fig. 2C). The resulting average GINI coefficients, defined as 2 × AUC −1 (where AUC represents area under the ROC curve), were 0.75 ± 0.04 and 0.63 ± 0.04 for Gaussian and Non-rigid data sets, respectively, with the restricted extraction. Average GINI coefficients with the unrestricted extraction were higher, with the respective coefficients being 0.87 ± 0.02 and 0.82 ± 0.02 for Gaussian and Non-rigid data sets. These quantitative metrics demonstrate that SCOUT is a robust classifier across different datasets (Fig. 2C).

### Comparisons of SCOUT with CNMF-E on in vivo single session recordings

After verifying the effectiveness of SCOUT on simulated data, we continued to examine the effects of introducing spatial constraints to neuron extraction in experimental *in vivo* recordings from hippocampal CA1, obtained from three separate mice (see **Materials and Methods**). Each recording consisted of approximately 7000 frames each and was extracted using both CNMF-E and SCOUT. While ground truth data was not available for these recordings, we developed a set of three criteria for classifying neurons as true discoveries. First, the spatial footprint was examined visually. Neurons with abnormal footprints were removed from consideration. Next, the fluorescence trace corresponding to each neuron was examined for irregularities, such as traces with non-zero baselines, or traces that exhibited localized activity that may be attributable to recording noise, or poor extraction quality. Finally, the remaining neurons were plotted on the correlation image (which shows local correlation between neighboring pixels, Supplementary Fig. 1C, Fig 2E **(1,2) top subpanels**), and neurons that appear to encompass spatially distinct regions of the correlation image were removed as false discoveries. Across all three recordings, an average of 24.3 ± 2.3% of neurons discovered by CNMF-E, were classified as false discoveries, while 9.0 ±2.1% of neurons discovered by SCOUT were classified as false discoveries (Fig. 2E).

We also tested SCOUT on three additional recordings, one each from the visual cortex, the hippocampal CA1, and the prefrontal cortex, having 4000-8000 frames each. CNMF-E detected an average of 15% more neurons than SCOUT, although after false discovery removal. The number of detected true discoveries was, on average, larger with SCOUT than with CNMF-E. In these additional datasets, we found an average of 24.4 ± 7.9% of neurons detected by CNMF-E were classified as false discoveries, while 8.1 ± 0.7% of neurons detected by SCOUT were classified as false discoveries.

### Comparisons of SCOUT and cellReg on simulated multi-session recordings

We next compare SCOUT with cellReg on multi-session simulated recordings, to empirically demonstrate that SCOUT tracks significantly more neurons than cellReg, with comparative or lower false discovery rates. Both methods were tested on three sets of simulated data: (1) a subset of 11 of the Gaussian recordings discussed earlier (splitting sessions every 2000 frames to obtain multi-session sets of sub-recordings for cell tracking) referred to as the Fixed dataset, (2) a set of 39 recordings consisting of 8000 frames each, split into 4 sub-recordings of 2000 frames, in which random non-rigid transformations are applied to each spatial footprint in each sub-recording, referred to as the Non-Rigid dataset, and (3) a simulated dataset in which no background noise was included, with each recording having 6000 frames with Gaussian simulated neuron footprints individually shifted every 3000 frames, creating two sub-recordings for each recording (shift for each footprint was independent and less than 30% average neuron width, recordings split every 3000 frames for cell tracking), referred to as the Shifted dataset (Fig. 3, Supplementary Fig. 2, **Materials and Methods**).

**Figure 3.**
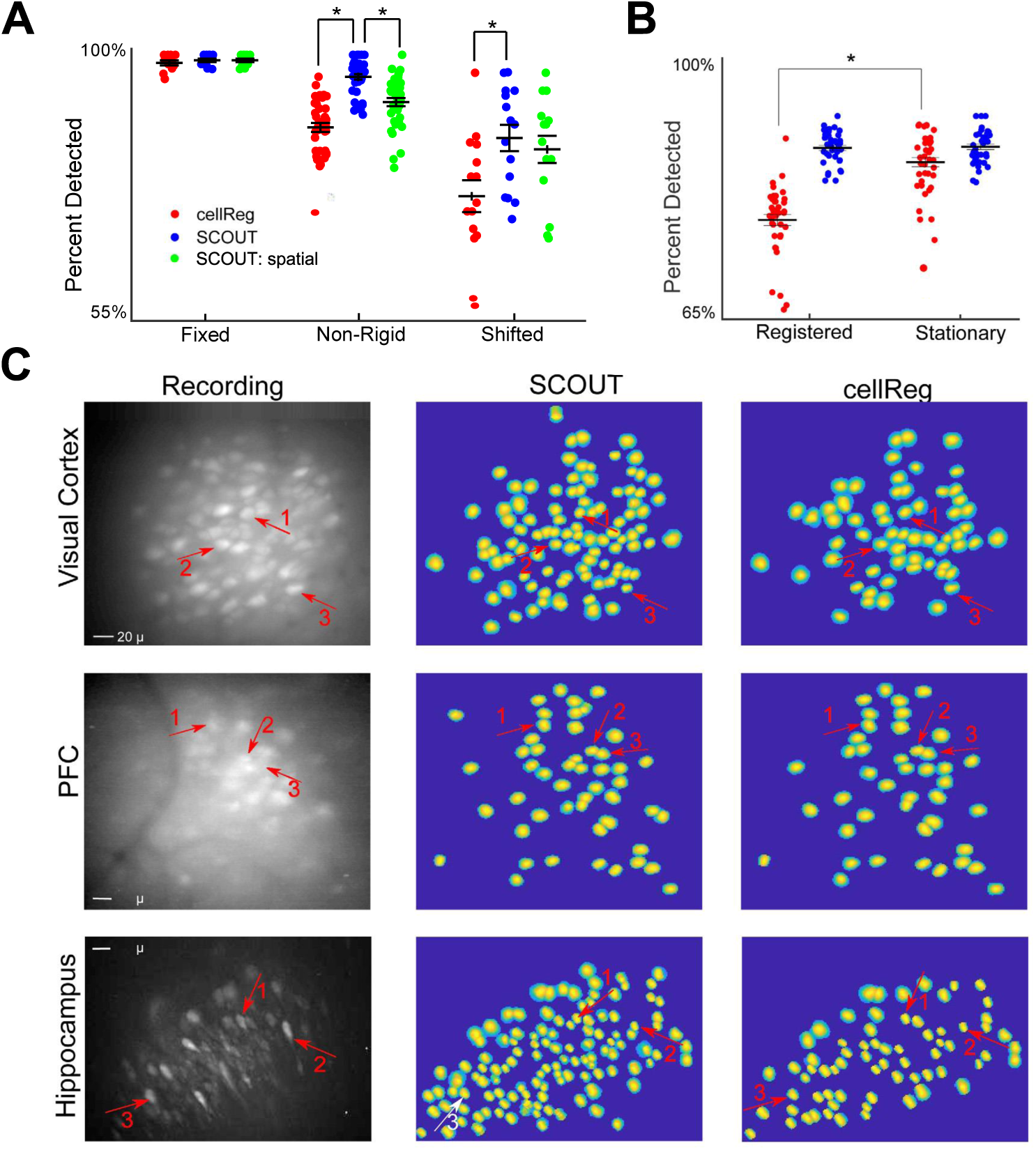
SCOUT enhances neuronal detection in multiple session recordings. **A:** We compare average percent detected rates of cell tracking using cellReg, SCOUT, and SCOUT spatial (which only uses spatial criterion), based on SCOUT extractions of each recording. This is calculated as the percentage of neurons tracked through an entire recording, divided by the total number of neurons that were extracted in each sub-recording. Asterisks indicate statistical significance calculated via t-test. Note that SCOUT tracks significantly more neurons on average than cellReg on each dataset. On the Non-rigid dataset, we see the importance of including temporal correlation. **B:** On the Non-rigid dataset with CNMF-E extractions, the PDR for cellReg is significantly less than the PDR for SCOUT when using session registration prior to cell tracking (Registered), compared with cell tracking without session registration (Stationary). **C:** Comparison of cell tracking methods on recordings taken from three distinct regions of the brain: the visual cortex, prefrontal cortex, and hippocampus. All three regions show increase in the number of neurons tracked by SCOUT over cellReg. For each region, 6-7 recordings were taken, each consisting of 4000-13000 frames. Sessions were extracted individually (including connecting sessions for SCOUT), with consequent cell tracking using both methods. Comparisons of tracked neurons from SCOUT (2^nd^ column) and cellReg (3rd column) show high spatial fidelity with those visually identifiable in the template (obtained via maximum projection of the recording, 1^st^ column). SCOUT identified significantly more neurons in both the visual cortex and the hippocampus.

Cell tracking for recordings in the Fixed dataset is a simple task, as no transformations are present between sessions. The Non-Rigid dataset simulates non-rigid spatial transformations that take place between sessions and is the most representative of true *in vivo* conditions. The Shifted dataset includes significant spatial translation between sessions, but with no non-rigid warping of spatial footprints. Cell tracking over this dataset is the most challenging task. No registration was performed prior to recording extraction. Connecting recordings were constructed using 1000-2950 frames from each consecutive pair of sub-recordings.

Simulated neurons in sub-recordings and connecting recordings were extracted via SCOUT and CNMF-E (except for the Shifted dataset in which the sub-recording extractions were virtually the same due to the lack of background signal), and cell tracking performance was compared between SCOUT, SCOUT: spatial (a version of scout using only spatial similarity measures), and cellReg with each dataset and extraction method. For a given recording, the output of cell tracking is a proposed chain of neurons, one from each sub-recording, and the associated concatenated neural signal, which should match one of the ground truth neurons.

As SCOUT requires both base recordings and connecting recordings, we cannot use the ground truth cell register. Instead, we used the sub-recording extractions to construct a cell register as follows: on each sub-recording, if a neuron extracted from that sub-recording had spatial correlation greater than 0.65, and temporal correlation greater than 0.8, with a ground truth neuron, the extracted neuron was identified with the ground truth neuron. Ground truth neurons identified in all sub-recordings were considered available. Statistics for cell tracking were then relayed in terms of PDR (the percentage of available neuron chains detected by the cell tracking method) and FDR (the percentage of detected neurons not corresponding to an available neuron). Here, we are distinguishing only between completely correct neuron chains, and all other detected neuron chains.

For each dataset and extraction method, we performed multiple cell tracking analyses with SCOUT, SCOUT: spatial, and cellReg, by varying parameters such as registration method and maximal distance between neighboring neurons (see **Materials and Methods** for parameter details). Presented results were obtained by choosing the cell tracking result that optimized the metric PDR/(1+FDR), a metric which penalizes high FDR, and rewards high PDR. Thus, presented results may be considered best case scenarios for each method. We found significant differentiation when comparing average PDR on all simulated datasets except the Fixed dataset (ANOVA Non-rigid: *p* = 1.1 × 10^-14^ (SCOUT extraction) *p* = 6.7 × 10^-8^ (CNMF-E extraction), Shifted: *p* =.015). Comparable FDR (i.e. lacking sufficient average difference to reject the null hypothesis) were detected across each dataset and method, except on the Non-rigid dataset, (ANOVA *p* < 3.1 × 10^-4^, for both CNMF-E and SCOUT extractions).

Investigating pairwise differences between SCOUT and cellReg, we find SCOUT detects significantly more neurons on the Non-Rigid and Shifted datasets (t-test Non-rigid: *p* < 1.3 × 10^-5^, Shifted: *p* = 1.7 × 10^-3^). SCOUT had comparable false discovery rates with cellReg (i.e. statistical comparisons did not reject the null hypothesis) on all datasets except Non-rigid, where SCOUT did significantly better (*p* = 2.7 × 10^-9^, SCOUT extraction only).

Comparing SCOUT and SCOUT: spatial yields significant differentiation in PDR only with the Non-rigid dataset (PDR: *p* < 3.3 × 10^-11^, FDR: *p* < 1.2 × 10^-8^). This is expected, as with the Fixed dataset, virtually all cells were found using both versions of SCOUT, and with the Shifted dataset, distances between identified cells were significant enough that extracted single neurons in connecting recordings failed to accurately account for signals on the overlap (Fig 3A, Supplementary Fig 2A, Table 2).

**Table 2:** Comparison of PDR and FDR across cellReg, and cell tracking with SCOUT, and SCOUT: spatial. The outcome maximizing PDR/(1+FDR) is highlighted in bold. Reported as mean ± standard error.

| Dataset | Extraction Method | CellReg |  | Tracking with SCOUT |  | Tracking with SCOUT: Spatial |  |
| --- | --- | --- | --- | --- | --- | --- | --- |
|  |  | PDR (%) | FDR (%) | PDR (%) | FDR (%) | PDR (%) | FDR (%) |
| Fixed | CNMF-E | 95.5 ± 0.7 | 3.4 ± 0.7 | **97.5 ± 0.7** | **2.1 ± 0.6** | 95.2 ± 1.0 | 3.6 ± 1.2 |
|  | SCOUT | 98.7 ± 0.2 | 2.6 ± 0.6 | 99.1 ± 0.3 | 3.3 ± 0.6 | **99.1 ± 0.7** | **3.2 ± 0.6** |
| Non-rigid | CNMF-E | 90.4 ± 1.3 | 9.8 ± 1.3 | **94.7 ± 0.8** | **8.8 ± 1.2** | 90.3 ± 0.9 | 12.8 ± 1.5 |
|  | SCOUT | 87.8 ± 1.4 | 10.6 ± 1.2 | **96.3 ± 0.9** | **6.2 ± 0.1** | 92.0 ± 1.3 | 10.1 ± 1.3 |
| Shifted | SCOUT | 76.1 ± 3.2 | 32.9 ± 2.6 | **86.0 ± 2.6** | **32.1 ± 2.9** | 84.0 ± 2.7 | 32.9 ± 2.9 |

One important caveat of using cellReg’s built-in session-registration methods appear to be significantly affected by false discoveries in the sub-recording extractions (Fig 3B). This is particularly relevant to the CNMF-E extractions, as average PDR of cellReg with its built-in registration was 82.1%. In comparison, without cell eg’s session-registration, the average PDR was significantly higher at 92.1% (*p* = 3.5 × 10^-11^), with significantly lower false discovery rates (*p* =.1.2 × 10^-5^). SCOUT did not result in such a decrease in PDR (94.5% vs 94.7%, registered vs stationary). The version of cellReg used by the authors did not include an option not to register sessions. The authors edited the code base to obtain the above results.

### Comparisons of SCOUT and cellReg on in vivo multi-session recordings

We continued to compare the results of SCOUT and cellReg on several *in vivo* recordings from the visual cortex, prefrontal cortex, and hippocampus of mice (Fig. 3C). We used 6-7 recordings (4000 –8500 frames each) from each region and compared the results of neuron tracking using SCOUT and cellReg. For both methods, parameters were varied to produce the maximal number of identified neurons. Connecting recordings consisted of 8000 frames (4000 from each pair of consecutive recordings). Visual identifications were used to eliminate probable false identifications between sessions (**Supp. Video 2**). On average, 11.7 ± 3.5% of neurons detected by cellReg were classified as false discoveries, while 4.6 ± 2.6% of neurons detected by SCOUT were identified as false discoveries. After false discovery removal, cellReg identified 62.0 ± 10.1 neurons in each set of recordings, while SCOUT identified 100 ± 9.5 neurons, consistently identifying more neurons than cellReg in each session.

Taken above together, SCOUT shows significant improvements over earlier methods including CNMF-E and cellReg in identifying neurons across sessions. Overall, SCOUT exhibited comparable or fewer false discoveries than cellReg, while consistently detecting more neurons, across simulated and *in vivo* recordings.

### Longitudinal analysis of contextual discrimination associated ensemble neuronal dynamics

Using SCOUT, we obtain improved population cell tracking of behavior-associated hippocampal neural ensemble dynamics at single-cell resolution for longitudinal analysis. We applied SCOUT to miniscope imaging data analysis of long-term contextual discrimination acquisition, extinction, and reinstatement (Fig. 4, Supplementary Fig. 1A). Mice were trained to acquire a context-specific fear response in one context (the stimulus context), which was paired with a mild footshock (0.25-0.5 mA), but not in a similar though visually distinguishable context (the control context) in which no footshock was administered. We collected both imaging and mouse behavioral data from the two contexts during learning, extinction, and relearning (reinstatement) stages of the experiment (Fig. 4A-E). In total, we tracked an average of 135 ± 21 neurons extracted per mouse across 5 mice, throughout the contextual discrimination experiment that lasted approximately two months, with between 36 and 44 recording sessions. Comparatively, cellReg detected an average of 77 ± 10 neurons, not including two mice for which cellReg returned no tracked neurons.

**Figure 4.**
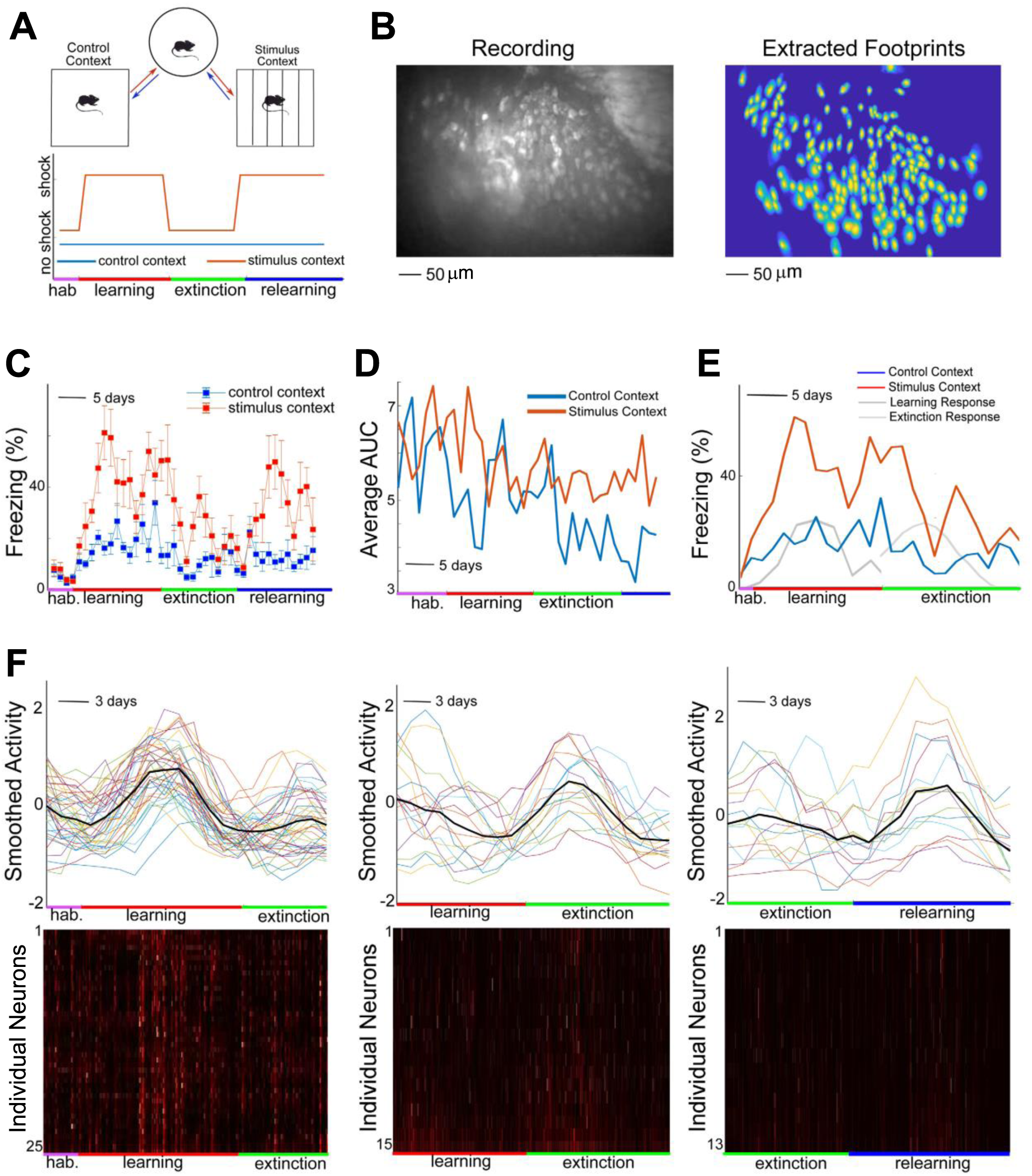
Longitudinal analysis of hippocampal ensemble activities during contextual discrimination experiments. **A**: (top) Visualization of the stimulus and control contexts, as well as the experimental process. Mice are placed in either the control or stimulus context for either 3 or 5 minutes (depending on context and experimental phase), then transitioned to a neutral context for twenty minutes, before being placed in the opposite context for another 3 or 5 minutes. (bottom) Visual representation of contextual discrimination procedure. After habituation (hab.), the mouse receives a single mild footshock in the stimulus context during two distinct periods (learning and relearning) separated by an extinction phase. In the stimulus context, the brief footshock is administered 3 minutes after being placed in the context. In the control context, no shock is administered. In panels c and d, freezing levels are measured for the 3-minute period after introduction to the context, and before the administration of footshock. **B:** Max projection image (left) and spatial footprints of extracted neurons (right) by SCOUT with data collected from a long-term contextual discrimination experiment for an individual mouse. For this mouse, 168 neurons were tracked across 38 sessions. **C:** During initial phases of training with footshock, mice exhibit behavioral generalization by increasing freezing in both contexts. After several days of training, mice exhibit contextual discrimination, which is evidenced by higher proportions of time freezing in the stimulus context as compared to the control context. During extinction, freezing decreases in both contexts, but shows a greater reduction in the stimulus context. Similar qualitative results occur during relearning (reinstatement) when footshock is applied again specifically in the stimulus context. Data shown is the mean time mice spent freezing in the first 3 minutes after placement in the respective context averaged for six mice (error bars indicate standard error of the mean). **D:** Each data point represents the area-under-curve (AUC) of the extracted neural calcium signals, averaged over all extracted neurons, for both contexts, for the specified day. Calculated AUC from the stimulus context does not include the time points after application of stimulus. Average neural activity indicates significant neural discrimination between contexts throughout the experiment. This effect is noted in three of the five mice. **E:** Freezing rates for a single mouse in stimulus (red) and control (blue) contexts, are compared with the mean cell activity across active neurons in the learning and extinction phases of the experiment (grey and black, respectively). Note the relative increase of neural activity during the peak of the acquisition and extinction phases, where changes in behavioral responses are most prominent. **F:** (top row) Each plotted point represents the smoothed daily area-under-curve (AUC) of the calcium signal trace for a single neuron. Here we plot daily AUC for a subset of neurons that exhibit increased activity during different stages of the contextual discrimination experiment. This panel demonstrates context-dependent neural activity changes observed across 3 of the mice (raw (unsmoothed) daily AUC is plotted in Supplementary Fig. 3B for reference). The data shows a clearly distinguishable increase in distributed activity with learning, extinction, and relearning. (bottom row) Raster plots of individual calcium signals for the corresponding neuron subsets show visually distinguishable increases in activity at the corresponding times above. Each row of the heatmap indicates the signal intensity for a single neuron, throughout the portion of the experiment indicated at the bottom. Signals were normalized to have the same maximum intensity for visualization purposes.

Context-dependent neural activity at the global level (calculated as the average area-under-curve (AUC) value across all neurons for each recording session) was exhibited in three of the mice (Fig. 4F). Two mice exhibited higher daily average neural activity in the stimulus context (not including footshock), compared to the control context, and one mouse exhibited increased daily average neural activity in the control context. For these mice, activity was higher in their preferred context 75.6 ± 4.3% of the time, a significant deviation from the average (*p* =.02, Welch’s t-test). The remaining experimental mice, average neural activity did not show significant preference for either context.

Additionally, we identified novel neural ensembles that exhibited a sustained (over 3-5 days) increase in activity followed by return to a baseline level. Across the three mice exhibiting global context preference, 32 ± 5.5% of all neurons exhibited stimulus-context dependent response changes in at least one context through the course of the experiment. At the learning stage, 16.7 ± 5.4% of the cells exhibited increased activity in either the stimulus context (prior to footshock) or control context within 1-5 days of imposition of shock, descending to baseline levels before the next phase of the experiment (Fig. 4F, left, top and bottom panels), while 13.2 ± 1.6% and 8.9 ± 2.3% exhibiting increased activity in the extinction and relearning phases (Fig. 4F, middle and right), starting 1-5 days after experimental phase change, respectively.

During learning, 7.7 ± 4.0% of neurons exhibiting increased activity (in at least one context) exhibited increased activity in both contexts, while only 4.8 ± 2.6 of neurons exhibiting increased activity during extinction exhibited increased activity in both contexts. A proportion of active neurons during extinction (<10%) exhibited increased activity across contexts or across learning phases.

## Discussion

In the present study, we have applied innovative concepts including spatial constraint priors and connected recording segments to develop a new method, SCOUT, for population single neuronal extraction and longitudinal tracking of their calcium responses across different recording sessions. As shown above, we demonstrate that SCOUT overcomes existing issues of earlier methods such as CNMF-E and cellReg for longitudinal analysis of miniscope-based recordings of neural activity at the single-cell level with significantly better performance.

Critically, SCOUT imposes spatial constraints on neuronal footprint extraction to control the number of false discoveries. This allows us to combine spatiotemporal correlation across recording sessions with a predictor-corrector methodology [Burden, et al. 1997] to develop a new method for the extraction of populations of single-cell neuronal data from long-term multi-session imaging experiments. This method presents measurably significant advancements, and our software implementation facilitates large-scale imaging data processing in automated and high throughput manners. SCOUT is highly parallelizable and has been tested on High Performance Clusters (HPCs) for extraction and cell tracking through large datasets. Considering the general-purpose nature of our filtering and signal recognition algorithms, we expect SCOUT to extend to multi-photon imaging data sets and applicable to other imaging modalities with some appropriate further modifications.

The introduction of a spatial constraint is shown not only to decrease the rate of false discoveries in the extraction process, but to increase the quality of the extracted calcium signal traces, in both fixed footprint and shifted footprint conditions. We create an elliptical model (spatial constraint) of neuronal footprints for 1p data and treat each spatial footprint as a discrete probability distribution. Each footprint is then compared with a baseline distribution using the Jensen-Shannon divergence metric that changes the sensitivity of the algorithm and allows the user to optimize the tradeoff between false discoveries and true positives. In our study, the addition of temporal similarity measures to spatial similarity, combined with a predictor-corrector methodology increases the number of detected neurons in longitudinally tracked neurons between sessions. This makes it possible to effectively track large numbers of neurons across potentially very large numbers of recordings. For proof of concept, we demonstrate this method utility across 60 days of consecutive recordings. In addition, probabilistic modeling is essential for our SCOUT method, as it provides a principled method for scoring pairs of neurons between recordings based on their spatiotemporal similarity, without imposing hard thresholds for any given similarity metric. We foresee that the new concepts and techniques used in SCOUT will improve many related cell registration and tracking methods.

Our validation using actual experimental data from multiple brain regions confirms the strengths of SCOUT over existing techniques. We have applied SCOUT to our longitudinal analysis of population neurons in a contextual discrimination to demonstrate its use for higher level neuronal transformations beyond simpler input/output functions. By tracking individual neurons throughout the experiment, we discovered evidence of population-wide differences that appear in relation to the application of shock stimulus as well as its removal. This provides evidence for pattern separation represented by evolving neural ensemble activities that are dependent on stimulus and context ([Czerniawski et al, 2014]). Our new methods enable robust longitudinal analysis of long-term imaging experiments, providing significant new insight into contextual discrimination-associated neural ensemble dynamics.

SCOUT code is publicly available on Github (https://github.com/kgj1234/SCOUT).

## Materials and Methods

### Simulated Recordings

Neuron footprints were simulated as 2-dimensional Gaussians, with diagonal covariance matrices Spatial footprint width was between 20 and 25 pixels. Spikes were simulated from a Bernoulli distribution with probability of spiking per timebin 0.01, and then convolved with a temporal kernel *g*(*t*) = *exp*(−*t/τ_d_*) − *exp*(−*t/τ_r_*), with fall time *τ_d_* = 6 timebins, and rise time *τ_r_* = 1 timebins. Local background spatial footprints were simulated as 2-D Gaussians, but with larger covariance entries than for the neuron spatial footprint. Blood vessel spatial footprints were simulated using a cubic function, that was convolved with a 2-D Gaussian (Gaussian width: 3 pixels). A random walk model was used to simulate temporal fluctuations of local background and blood vessels. 23 background sources were used throughout all simulated experiments, except for the Shift dataset, in which no background sources were present.

Three sets of recordings were simulated for testing purposes. The Fixed footprint simulated dataset consisted of 14 recordings with 2000-10000 frames each, with a 256 × 256-pixel FOV. Each recording was simulated using 50-200 neurons. We used two non-rigid footprint recording datasets, one for cell extraction consisting of 40 recordings having at least 2000 frames, and one for cell tracking consisting of 39 footprint recordings having at least 2000 frames. Each simulated spatial footprint was transformed with a non-rigid transformation for the cell extraction comparison and transformed every 2000 frames in the cell tracking comparison. Individually shifted footprint recordings consisted of 14 recordings with 6000 frames each, with a 100 × 100-pixel FOV. Each recording was simulated using 50 neurons. The individual spatial footprints were shifted independently by between 5 and 7 pixels in the latter 3000 frames (creating two sub-recordings of length 3000 frames in which spatial footprints are consistent, for each recording).

### Animal experiments and miniscope recordings

All animal experiments were conducted according to the National Institutes of Health guidelines for animal care and use and were approved by the Institutional Animal Care and Use Committee and the Institutional Biosafety Committee of the University of California, Irvine.

#### Viral injections

To perform viral injections, mice were anesthetized under 1.5% isoflurane for 10 minutes with a 0.8 L/min oxygen flow rate using a bench top unit (HME1-9, Highland Medical Equipment). Carprofen and buprenorphine analgesia were administered preoperatively. Mice were then placed into a stereotaxic unit for mice (Leica Angle Two) with their heads secured and received continuous 1% isoflurane anesthesia. Core body temperature was maintained at 37.5°C using a feedback heating system. Eyes were coated with a thin layer of ophthalmic ointment to prevent desiccation. The skull was then cleaned with iodine and 70% ethanol. A small incision was made on the scalp and the skin was opened to expose the skull and the landmarks of bregma and lambda.

A three-axis micromanipulator guided by a digital atlas was used to determine the position of bregma and lambda. Using the micromanipulator software, the injection site was calculated relative to bregma and lambda, using computerized coordinates in the digital atlas. The injection coordinated targeting bV1 are anteroposterior (AP) −3.4 mm, mediolateral (ML) 2.75 mm, and dorsoventral (DV) −1.20 mm (all values are relative to the bregma). At the injection site, a small drill hole was made in the skull, exposing the pia surface. Then, a glass pipette (tip diameter, ~20-30 µM) loaded with virus, was lowered into the brain to the appropriate depth and coordinates. Virus was pulsed into the brain at a rate of 20-30 nL/min with 10ms pulse duration using a Picospritzer (General Valve, Hollis, NH). Backflow of virus was prevented by allowing the pipette to remain in the brain for 5 min after the completion of the injection. Upon withdrawal of the injection pipette, the mouse was removed from the stereotaxic frame, and the scalp was closed with tissue adhesive (3M Vetbond, St. Paul, MN). Mice were injected with 5mg/kg Carprofen to mitigate pain and inflammation. Mice were then taken back and recovered in their home cages.

#### Miniscope imaging preparation, GRIN lens implantation, and baseplate placement

At two weeks after AAV1-CaMKII-GCaMP6f injection, a gradient refractive index (GRIN) lens was implanted at the injection site in CA1. A.8 mm-diameter circular craniotomy was centered at the coordinates (AP −2.30 mm and ML +1.75 mm relative to bregma). ACSF was repeatedly applied to the exposed tissue; the cortex directly below the craniotomy was aspirated with a 27-gauge blunt syringe needle attached to a vacuum pump. The unilateral cortical aspiration might affect part of the anteromedial visual area determined using Allen Brain Atlas (http://www.brain-map.org/), but the procedure left the primary visual area intact. The GRIN lens (0.25 pitch, 0.55 A,.8 mm diameter and 4.3 mm in length, Edmund Optics) was slowly lowered with a stereotaxic arm to CA1 with a depth of −1.60 mm relative to the bregma. Next, a skull screw was used to anchor the GRIN lens to the skull. Both the GRIN lens and skull screw were fixed with cyanoacrylate and dental cement. Kwik-Sil (World Precision Instruments) was used to cover the lens. Two weeks later, a small aluminum baseplate was cemented onto the animal’s head atop the existing dental cement. A miniscope was fitted into the baseplate and locked in a position so that the field of view was in focus to visualize GCaMP6f expressing neurons and visible landmarks, such as blood vessels.

Please refer to http://www.miniscope.org/ for technical details of our custom-constructed miniscopes. The head-mounted scope has a mass of about 3 grams and uses a single, flexible coaxial cable to carry power, control signals, and imaging data to custom open source Data Acquisition (DAQ) hardware and software. Under our experimental conditions, the miniscope has a 700 μm × 450 μm field of view with a resolution of 752 pixels × 480 pixels (~ μm per pixel). The electronics packaged the data to comply with the USB video class (UVC) protocol and then transmitted the data over Super Speed USB to a PC running custom DAQ software. The DAQ software was written in C++ and uses Open Computer Vision (OpenCV) libraries for image acquisition. Images were acquired at ~30 frames per second (fps) and recorded to uncompressed .avi files. The AQ software simultaneously recorded animal’s behavior through a high definition webcam (Logitech) at ~30 fps, with time stamping both video streams for offline alignment.

#### Contextual Discrimination

Mice were trained to differentiate between two similar, but visually distinct, square open field environments; miniscope imaging of hippocampal CA1 excitatory neurons was simultaneously conducted in behaving mice during the tasks (Fig. 4A-B). Mice were habituated for a two-week period, in which they allowed to free explore each environmental context daily. At the end of this habituation phase, context discrimination training started by introducing a mild foot-shock stimulus after 3 minutes in the stimulus context but not in the control context. The mice learned to freeze as a contextual discrimination response in anticipation of the stimulus. Subsequently, a two-week extinction phase in which no shock was applied, led to reduction in discrimination and freezing behavior. We then reinstated the stimulus to study neural response during reacquisition of the discrimination behavioral response.

Each day, individual mice were introduced to a random context (either control context or stimulus context), followed by 20 minutes in a neutral context, after which the mice were introduced to the remaining context. Recordings were taken in both contexts. Mice spent 3 minutes in the control context, and 5 minutes in the stimulus context. The recordings from the stimulus context were split into a 3-minute baseline recording, and a 2-minute stimulus recording, in which a 1s 0.25-0.5 footshock was applied, 30 seconds into the recording. During the habituation and extinction phases, no stimulus was delivered. Cages were thoroughly cleaned between sessions involving different mice (Fig 4B; Supplementary Fig. 3A-B). Mouse freezing behavior, as evidenced by a lack of movement except that necessary for respiration, was manually measured offline from video recordings of the session.

#### Evidence for Localization of Cell Response after Introduction or Removal of Stimulus

The initial K-means clustering applied to daily neural activity (as measured by AUC) implied the existence of a sustained activity increase occurring within 1-5 days of the initiation of the learning, extinction, and relearning phases in a significant subset of neurons, in both contexts, for nearly all mice. In order to test this theory, and demonstrate that directly after introduction and removal of stimulus were the only three probable stages in the learning process where a significant, sustained increase in activity occurs among a large subset of extracted neurons, we created a comparison template of length 11 days, with behavior similar to the detected behavior (namely increase in activity over four days, followed by 3 days holding steady, followed by decrease in activity). The number of neurons exhibiting similar activity patterns (based on a correlation of 0.65 with the template) starting at each day of the trial was counted, and significantly more neurons followed this activity pattern within two days of the imposition or removal of stimulus, than the average at other time periods in the experiment, indicating that the responses were stimulus dependent. Modifying the template in terms of activity rate increase and sustained activity intensity did not significantly alter the results, though decreasing the template length resulted in the detection of additional neurons with characteristic activity starting at each day, diluting the uniqueness of the effect.

### Preprocessing recordings

*In vivo* recordings were preprocessed using NoRMCorre image registration for motion correction [Pnevmatikakis and Giovannucci, 2017]. For experiments taking place over more than one recording, alignment between sessions was performed either manually, by using max projections in imageJ [Schindelin et al., 2012], or automatically using image registration libraries created for Matlab [Forsberg, 2015]. SCOUT provides an interface for automatic image registration, as well as manual feature selection-based registration.

### Optical recording extraction algorithms

One class of methods for signal extraction involves semi-manual ROI selection. Such methods include manual ROI selection of individual neuron footprints, and subsequent deconvolution of the neural trace, as well as methods such as convolutional neural networks (CNNs) which use a corpus of identified footprints to train a neural network to identify footprints in future experiments [Apthorpe et al., 2016], followed by a second step in which temporal fluorescence traces are extracted based on the proposed footprints. However, such methods become computationally intractable when considering large cell population and become less accurate when considering neurons exhibiting strong spatial overlaps between footprints.

Another class of methods involves automated ROI construction, where both fluorescent traces, and spatial footprints are extracted simultaneously. The simplest such example is PCA/ICA [Mukamel et al., 2009], in which PCA and ICA are successively used to isolate and extract spatial footprints and spike trains from optical recordings. These methods rely on linear demixing and can produce significant error when neuron footprints exhibit strong spatial overlaps [Pnevmatikakis et al., 2016].

The most recent major advance in 1-photon optical recording extraction (as far as the authors are aware) is CNMF-E [Zhou et al., 2018]. As this is the primary method adapted in this paper, we will briefly describe the algorithm.

Given a recording, let *d* represent the number of pixels in the field of view, *T* the number of frames observed, and *K*, the number of neurons in the field of view. Then let *Y* ϵ 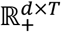 represent the initial calcium fluorescence recording; let *A* ϵ 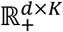, the spatial footprints of the neurons, with each column representing the footprint of a single neuron; let the rows of *C* ϵ 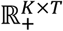 represent the fluorescent signal of each neuron at each frame; and let *B* ϵ 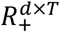 represent the background fluctuation. The goal is to find *A,B,C* such that |׀*Y* – (*AC* + *B*)׀|*_F_* is minimized, which can be interpreted as determining the optimal spatial footprints, fluorescence traces, and background noise, in order to reconstruct the recording.

The *i*th row of *C* is represented as an autoregressive process, where 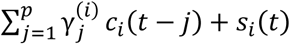 and *s*_*i*_(*t*) represents the number of spikes fired by the *i*-th neuron in the *t*th frame, and *S*, the matrix of spikes, is constrained to be sparse. The footprint matrix *A* is also constrained to be sparse, and *B* is constrained to be a nonnegative matrix decomposable as *B* = *B*^*f*^ + *B*^*c*^ Where *B*^*c*^ models the constant baseline background, and *B*^*f*^ models fluctuating background activity. Initialization for neuron centers uses a greedy algorithm, such that a proposed pixel satisfies two criteria: a minimum threshold on peak-to-noise ratio (calculated as peak signal strength divided the standard deviation of the noise), and a sufficiently high temporal local correlation (implying strong similarities in temporal signal for pixels surrounding the proposed center) ([Smith et al, 2010], Supplementary Fig. 1B-D). Initialization of variables *C* and *B*, as well as updates for the background *B* are discussed in the original paper. [Zhou et al., 2018].

### False discovery removal via SCOUT

Regarding false discovery removal, after each iteration of the extraction method, an initial pre-processing begins in which proposed spatial footprints are thresholded based on maximum pixel intensity, removing low intensity (<10% of maximum intensity) pixels. Each footprint is normalized so that the sum of pixel intensities in each footprint equals 1, allowing us to view each spatial footprint as a discrete probability distribution. Each footprint is then compared with a baseline distribution, using Jensen-Shannon (JS) divergence as a metric [Kullback. S, 1997] (Supplementary Fig. 1E-F). Subsequently, the footprints with similarity exceeding the baseline of a specified JS threshold are removed, and the remaining footprints are updated either by averaging with the baseline, or by setting the pixel intensities of all points not in the support of the baseline to 0. Using an iterative process employed by the CNMF-E, the spatiotemporal traces corresponding to each footprint are then updated by removing any non-zero intensity levels not in the support of the comparison footprint, after which the remaining intensity values are rescaled to their original magnitude (Fig. 1C-D). Note that varying the JS threshold changes the sensitivity of the algorithm (Supplementary Fig. 1D-E, Fig. 2C), allowing the user to optimize the tradeoff between false discoveries and true positives. Baseline neurons footprints are sampled from a user determined probability distribution, with parameters sampled from the proposed distribution.

While the construction of the baseline is customizable, we consider several options for 1p data (Supplementary Fig. 1E). One option is a Gaussian model. Mean and covariance are sampled from the normalized, thresholded footprint *P*, and used to construct a comparison footprint *Q*. We found that Gaussian models overestimated the rate of decrease in signal intensity when moving from the center of the proposed neuron to its boundary. Another is an elliptical comparison created by calculating the centroid from the footprint *P* and the rate of signal strength intensity decrease along the major axis of the footprint. The intensity values of the footprint are interpolated along the major axis using a fractional polynomial model, and the fractional polynomial is rotated around the centroid of the footprint, linearly scaled so as to decrease to the width of the minor axis after a rotation of 90°, to create an elliptical model for the neurons spatial footprint. This is the method used in the experimental results discussed in the main body, as it appeared to show greater differentiation between true and false discoveries, than the Gaussian model.

### Comparison of spatial filter with alternative methods

While machine learning algorithms have previously been used to classify false discoveries derived from neural extraction, there are two significant issues for their usage.

First, they frequently require additional training on a labeled dataset. For example, we tested both CaImAn (which only has a network trained on 2p data), and an AutoML curation algorithm [Tran et. al., 2020], and neither gave decent results without retraining (treating them as binary classifiers, the ROC curve had AUC less than ½). The AutoML algorithm advised a training time of two days and required a specific data input format that did not easily generalize to different magnifications.

Second, both networks required significant additional software to run. The AutoML algorithm was implemented only in python, making it difficult to include in MATLAB pipelines, and CaImAn requires the neural network toolbox, as well as some additional open source software. Our spatial filter runs in base MATLAB.

One benefit of using a spatial filter over a machine learning algorithm is in interpretability. For each neuron, we can identify the closest member in an associated probability distribution and measure the distance between them. This allows us to precisely identify why the algorithm classifies each entry as positive or negative.

### Cell Tracking via SCOUT

Given two registered recordings, we construct a connecting segment between the two recordings, consisting of frames from the end of the first recording and the beginning of the second, to form a recording overlapping portions of the initial recordings (Fig. 1C). Next, we extract the neural activity from all three recordings. The connecting recording, though typically having lower extraction quality, can be used to identify temporal traces between sessions, via correlation on the overlap.

After these preliminary steps, we perform a predictor step, in which initial identification probabilities are assigned between neurons in the two initial recordings, using spatial and temporal criteria. Spatial footprint similarity between neurons can be calculated using metrics such as centroid similarity, spatial overlap, and JS. Temporal metrics, such as correlation on the overlap, signal-to-noise ratio, and decay rate, can be used to provide additional differentiation for spatially similar neurons.

Methods such as mixture models are used to assign probabilities to the identification based on the spatial and temporal correlation between identified neurons in the initial sessions for each chosen metric, similar to cellReg [Sheintuch et al., 2017]. The probabilities are weighted and used to create a single identification probability between each neuron pair, creating an ensemble probabilistic classifier.

Next, we apply a stochastic process as follows to adjust identification probabilities, so that the greatest number of neurons would be detected. Initially, we construct a transition matrix consisting of identification probabilities between all possible neuron pairs between sessions. This is divided into connected components, consisting of interconnected neuron identifications. For each such connected component, we initialize the resulting neuron identifications using a greedy algorithm (i.e., we identify the strongest identification probability, and identify the corresponding neurons between sessions, then delete all probabilities associated with the identified neurons, and continue the process till all possible neurons are identified). We assign a negative penalty probability to any unidentified neurons and calculate the sum of probabilities for this identification. Finally, we perform several hundred iterations in which we randomly delete identification probabilities between neurons, to maximize the total sum of probabilities for this component. Deletion of certain identification probabilities has been shown to increase the total number of detected identifications using the greedy algorithm, decreasing connecting probabilities between some neurons, but increasing the overall sum of probabilities for the component. This showed the greatest effect on the individually shifted data, as occasionally, the nearest neuron (using the weighted similarity metrics) was not the correct identification.

Subsequently, probabilities are normalized. For a given neuron *N*_1_, if multiple identifications to neurons in the second recording exist, labeled 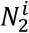, with associated probabilities 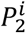, we calculate the new identification probability as 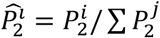, where the sum is taken over all possible identification probabilities for that neuron.

Using pairwise identifications for successive recordings, we track neurons through multiple sets of recordings by creating a connecting segment of recordings between each consecutive pair of recordings followed by the application the previous method to obtain probabilities for each pair of recordings, constructing chains of identified neurons across the recordings.

Next, a corrector step is used to eliminate neurons identified in multiple chains. Each neuron chain is assigned an occurrence probability, based on the similarity of consecutive neurons. If more than one chain contains a given neuron from a recording, the chain that is most probable is accepted, with the duplicated elements in the remaining chain deleted. Partial chains are merged to create new chains, and the probability of the new chains are calculated. The process continues until no possible neuron chains are left. All chains with probability below a user set probability threshold (typically between 0.3 and 0.7) are removed from consideration (Fig. 1D).

To enhance identification and tracking of neurons between consecutive recordings, we may use spatiotemporal similarities to identify remaining neurons across sessions that are not temporally adjacent (i.e. at least two sessions apart in our ordering). While temporal correlation on overlap cannot be used in this case, as extracting this many connecting recordings would be prohibitively expensive, considering identifications across all pairs of recordings allows us to detect neurons that may be inactive in some sessions. However, this requires probability assignments across all pairs of recordings, requiring significant additional computations (O(n^2^) in the number of recordings). Occurrence probabilities are calculated using pairwise scores between each neuron in the chain. Using spatiotemporal similarity between all chain members typically results in a significant increase in the number of detected neurons, as well as cell tracking accuracy (assuming accurate initial registration of sessions), though it requires more time and computational resources. We apply this approach in the experimental results discussed in the paper.

### Calculation of temporal correlation across sessions

Given two preprocessed optical recordings *R_1_* and *R_2_*, we construct a connecting recording *R_c_* by concatenating the last *n* frames the first recording, with the first *n* frames of the second, where *n* is some number less than the number of frames in *R_1_* and *R_2_*. Next, we extract spatial and fluorescence traces from *R_1_*, *R_2_*, and *R_c_.* At this point, spatial overlap, and correlation on the overlapping frames, are used to track neurons through multiple recordings, as follows.

Given *N*_1_, a neuron from *R_i_*, and *N*_2_, a neuron from *R*_2_, we start by setting a maximal distance threshold *m*, that defines neighboring neurons. If the distance between the centroids *N*_1_ and *N*_2_ exceeds *m, N*_1_ and *N*_2_ would not be considered neighbors. Only neighboring neurons can be identified as the same between sessions. Next, given a similarity metric, we calculate the distance between *N*_1_ and *N*_2_ for every set of neighbors *N*_1_ from *R_i_*, and *N*_2_ from *R_2_*. Examples of spatial similarity metrics include centroid distance, overlap, and JS divergence.

For temporal correlation similarity, a similarity score is obtained for each neighboring neuron pair (*N*_1_ and *N*_2_) in the two recording sessions, by ranging over the full set of neighboring neurons (*N*_*c*_) in the connecting recording (i.e. across the set of *N*_*c*_ coming from *R_c_* such that *N*_1_ is a neighbor to *N*_*c*_, and *N*_*c*_ is a neighbor to *N*_2_). The choice *N*_*c*_ that maximizes the average of the correlation distance between *N*_1_ and *N*_*c*_, and *N*_*c*_ and *N*_2_, is considered the connecting neuron, and the distance between *N*_1_ and *N*_2_ can be considered as the mean of the maximal correlation across choice of connecting neuron *N*_*c*_. Use of temporal correlation is not necessary for application of the algorithm, but there is some indication that using the correlation metric increases the percentage of correctly determined identifications, particularly on the individually shifted data (*p* = 0.08). For long term experiments, concatenation, and extraction of multiple recordings, followed by cell tracking via SCOUT provided the best results, and replaced the requirement for the separate extraction of *R_c_* by using overlapping concatenated batches of recordings.

### Spatial similarity measures for calculating neuron similarity across sessions

Currently, three methods for spatial similarity are included with SCOUT: centroid distance, spatial overlap, and Jensen-Shannon divergence. Centroids of neuron spatial footprints are calculated using the usual formulae 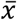 = Σ_*i,j*_ *X*_*i*_ *a*_*i,j*_, 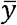 = Σ_*i,j*_ *y*_*j*_ *a*_*i,j*_ where *i,j* range across the number of pixels in the field of view, in the horizontal and vertical directions respectively, and *a*_*i,j*_ is the footprint intensity at the *ith* horizontal pixel, and the *jth* vertical pixel. Centroid distance between to footprints is calculated as the Euclidean distance between their centroids. Spatial overlap between footprints *a, b* is calculated as 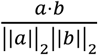,where *a* and *b*, are binarized column vectors representing whether each footprint has positive pixel intensity. Jensen-Shannon divergence between two (normalized) footprints *P,Q*, is calculated as 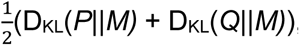, where 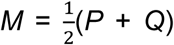, and D_KL_ is the Kullback-Liebler divergence: D_KL_(*P*||*Q*)=E(log[d*P/*d*Q*]), where d*P/dQ* is the radon-nikodym derivative of *P* with respect to *Q*.

### Temporal similarity measures for calculating neuron similarity across sessions

In addition to temporal correlation on connecting recordings, several additional temporal similarity measures can be deduced from properties of the fluorescence traces of each neuron. SCOUT has implemented temporal similarities based on signal-to-noise ratio (calculated as the average of the square of the signal strength, over the standard deviation of the estimated noise near the associated spatial footprint for each neuron), and the fluorescence trace decay rate for each neuron. Frequently, such similarities are preserved across recordings, and can be used to distinguish between possible identifications.

### Assigning Identification Probabilities with SCOUT

To assign probability scores between sessions for a given metric, we detail two approaches. First, we can simply assign the percentile as the probability score for each metric. If the distance between *N*_1_ and *N*_2_ for a given a metric, is less than *p*% of distances between all possible neighbor pairs, then *p* is the percentile assigned to the pairing. This method has several drawbacks. First, it is sensitive to the choice of maximum distance parameter. If the parameter governing the maximum distance between neighbors is increased, the probability assigned to any neighboring pair will increase. Second, when few neuron pairs exist, similarity metric values can accumulate near 0, so that even relatively small metric values can be associated to low probabilities. To avoid this problem, the distribution of metric values is approximated via kernel density estimation before the percentiles are calculated.

Another paradigm is to assume that for each metric, the distances between neighboring pairs come from a mixture of distributions: a distribution of distances corresponding the neurons that should be identified between sessions, and a set of neighbors that are distinct. Before fitting the mixture of distributions, a probability density function is constructed, by applying kernel density estimation to the normalized histogram of distances, using reflected boundaries near theoretical maximum and minimum values (such as 0 or 1 for correlation metrics). Next, we construct a model consisting of the weighted sum of two probability distribution functions, which is then fit to the approximated pdf, using nonlinear regression (Matlab nlinfit). We have implemented three cluster mixture models using Gaussian-Gaussian, Gaussian-Exponential, and Gaussian-Log Normal distributions [Everitt, 2014]. The default behavior is to approximate the pdf with each mixture model and choose the model that best approximates the pdf.

Given a mixture model consisting of a weight *w*, a model for identified neurons between sessions, *f*, and a model for unidentified neurons between sessions, *g*, the mixture model approximates the probability distribution function *h*, obtained via kernel density estimation from the initial distribution of distances, as *h(x)* = *wf(x)* + (1 - *w*)*g*(*x*). Given a proposed distance *x*, the probability that *x* is sampled from the distribution with pdf *f*, is given by 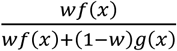 using Bayes theorem.

Another probabilistic clustering algorithm, soft K-means clustering [Dunn, 1973], an adaptation of K-means in which data points are assigned identification probabilities for each cluster, and a “fuzzifier” is introduced to govern the spread of identifications probabilities, adjusting the crispness of the clusters (Fig 1D). This algorithm frequently identified the most neurons, but with a higher false discovery rate.

### Probability Assignment for Neuron Chains in SCOUT

SCOUT provides several options for assigning probabilities to neuron chains. If only consecutive recording sessions are scored, then average (or minimum) probability between sessions are used to assign probability scores for each chain. When probability scores between all recording sessions are used, empirical results suggest a two-step method: first, a probability threshold is assigned, and occurrence probability is calculated as the minimum number of neurons any given neuron in the chain is connected to with probability higher than the probability threshold. Then, average chain probabilities across neuron pairs in the chain is used as a tiebreaker if multiple chains with the same connectivity scores are found.

### Long-term cell tracking with SCOUT

For long term cell tracking, we propose a combination of concatenation and cell tracking. In this methodology, recordings are concatenated into batches of uniform length, with overlapping portions of each batch used to calculate spatiotemporal similarity. For the contextual discrimination experiment, batches were composed of eight sessions, with an overlap of two sessions between batches. This method decreases the number of connecting recordings required. This method requires spatial footprint stability over each batch.

### Algorithm Parameter Settings

#### Cell Extraction

CNMF-E parameters were set as min_pnr = 5, min_corr = 0.8 (0.1 for unrestricted initialization), merge_thr = [1,1,-1] ([.65,.7,-1] on *in vivo* recordings), and dmin=0 ([1.5, 15] on *in vivo* recordings). All other parameters were left as defaults.

### Cell Tracking

For cell tracking via cellReg on the simulated recordings, we set p_same_threshold = 0.5, and performed 18 total cell tracking procedures, with varied parameters. maximal_distance (maximal distance between neighbors) varied between 10 and 50 by increments of 5.

For SCOUT, corr_thresh = 0.6, probability_assignment_method=Kmeans, chain_prob = 0.5, min_prob = 0.4. We performed 18 total cell tracking procedures with varied parameters. max_dist (maximal distance between neighbors) varied between 10 and 50 by increments of 5.

On *in vivo* recordings, the same parameters were used for cell tracking, but SCOUT was performed only once, with max_dist set to 45, and registration_method set to non-rigid, session registration for cellReg was set to non-rigid.

## Author contributions

X.X., K.G.J., J.G. and Q.N. conceived this work; K.G.J., S.F.G., Z.Y., R.C. performed the experiments; K.G.J. prepared illustrations; K.G.J., X.X., Q.N., T.C.H. wrote and edited the paper with the comments and inputs form all other authors.

## Acknowledgment

This work is supported in part by a BRAIN Initiative Grant (NS104897) from the US National Institutes of Health, an NSF grant DMS1763272 (Q.N.) and a Simons Foundation grant (594598, QN).

## Code Availability

Code is publicly available on Github (https://github.com/kgj1234/SCOUT).

---

**Supplementary Figure 1:**
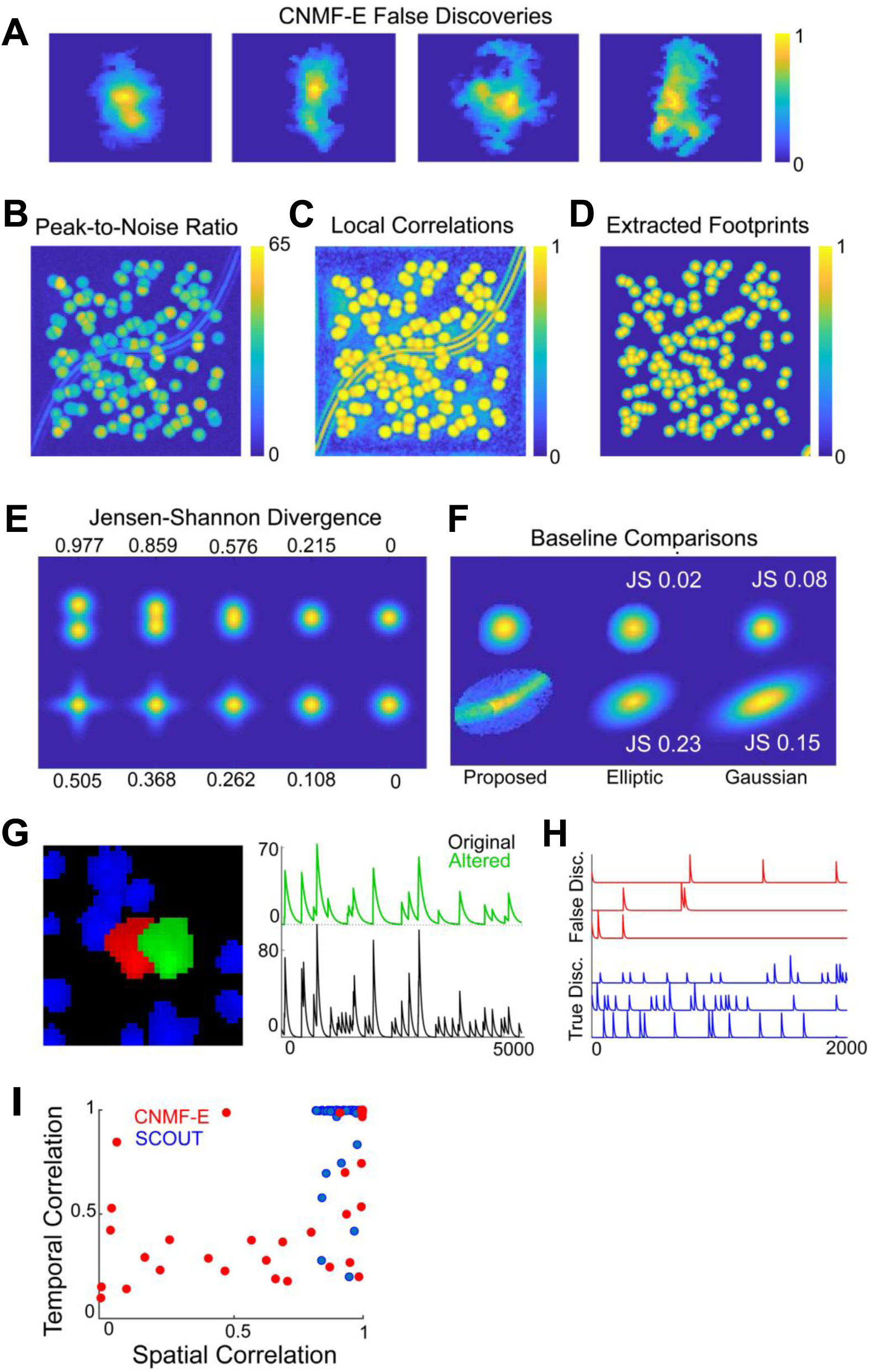
**A**: Examples of false discoveries detected by CNMF-E on *in vivo* recordings of the CA1 hippocampus. **B-D:** The local correlation image (**C**) and peak-to-noise ratios (**B**) are used as initialization points for neurons in CNMF-E and SCOUT. Extracted footprints (using SCOUT, **D**) show strong fidelity to the initialization values, and filter out most of the background activity. **E:** Examples of Jensen-Shannon divergence between Gaussian probability distributions which are (1^st^ row) gradually translated apart or (2^nd^ row) stretched in orthogonal directions. **F:** A true (top row) and false (bottom row) footprint discovery is shown compared with the elliptic comparison (2^nd^ column) and a Gaussian comparison (3^rd^ column). The elliptic comparison shows a stronger differentiation between the false and true discovery. **G:** Here we consider the effect adding an additional neuron (red), when extracting neural signals from a true neuron (green). On the right, we see a significantly altered neural signal (green) is reported than originally (black), when a spurious neuron is included during signal extraction. This illustrates the importance of removing false discoveries before the final signal update. **H**: We consider the extracted neural signals of several false discoveries obtained by CNMF-E on a recording in the Non-rigid dataset and compare with correctly detected neurons in the same dataset. On average, significantly fewer calcium signal events are detected in the false discoveries (top traces, red), as such signals are generated by background noise. **I**: Here we consider a correlation plot of extracted neurons, from one of the recordings in the Fixed dataset. Each point represents a neuron, with coordinates representing the maximal spatial and temporal similarities with ground truth neurons. The plot indicates that SCOUT has fewer false discoveries than CNMF-E. The footprint update procedure of CNMF-E can cause lower spatial correlation due to trimming of low intensity pixels. This effect can be adjusted by the user.

**Supplementary Figure 2:**
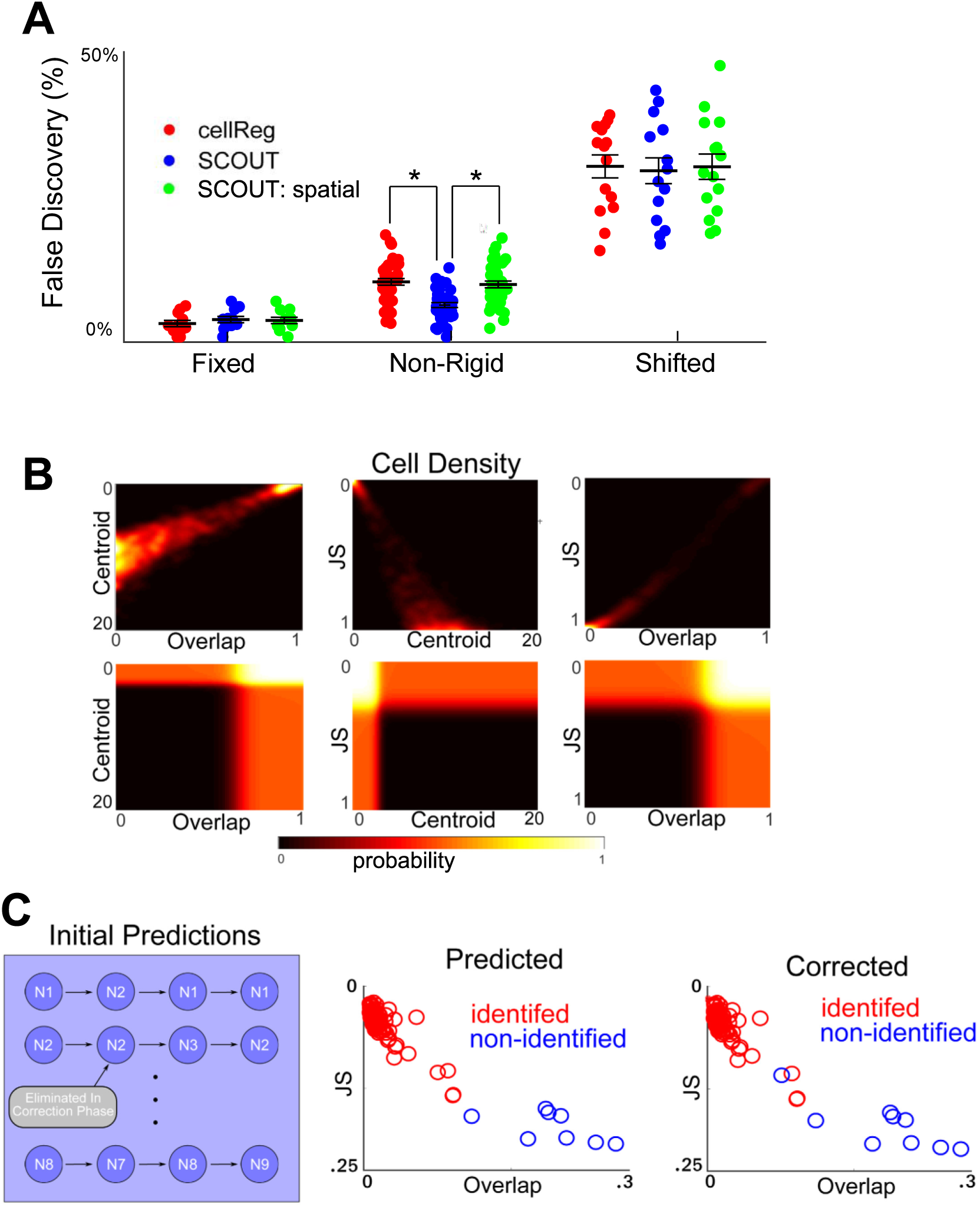
**A:** We consider the corresponding false discovery rates to cell tracking discussed in Figure 3A. In general, comparable false discovery rates are seen across all methods, though SCOUT has a significantly lower false discovery rate on the Non-rigid dataset. **B:** Top row figures show the density of neighboring cells using various distance metrics. Using Jensen-Shannon divergence, a clearer delineation between identified neighbors and non-identified neighbors between sessions is seen, viewed as low density in the middle regions of the plot, in both the second and third column. Bottom row figures show the probability density function created by applying a Gaussian mixture model to each density figure. **C:** (top) Initial probabilities between neuron pairs are used to create neuron chains across multiple recordings. This is followed by deletion of duplicate neurons based on which chains are identified as most probable. (middle and bottom) Plotting potential identified neighbors between sessions using their JS divergence and overlap distance demonstrates the predictor-corrector nature of the algorithm. Using simulated data an initial prediction is made, using a Gaussian mixture model, with a 0.5 cutoff for acceptance of probability. Following this, a correction is applied in which neurons in the initial session identified with more than one neuron in the secondary session, are eliminated based on aggregate probability, in this case, removing a single false discovery, forming a non-linear decision boundary. The corrected result identified all neurons in the recording, with no false discoveries.

**Supplementary Figure 3:**
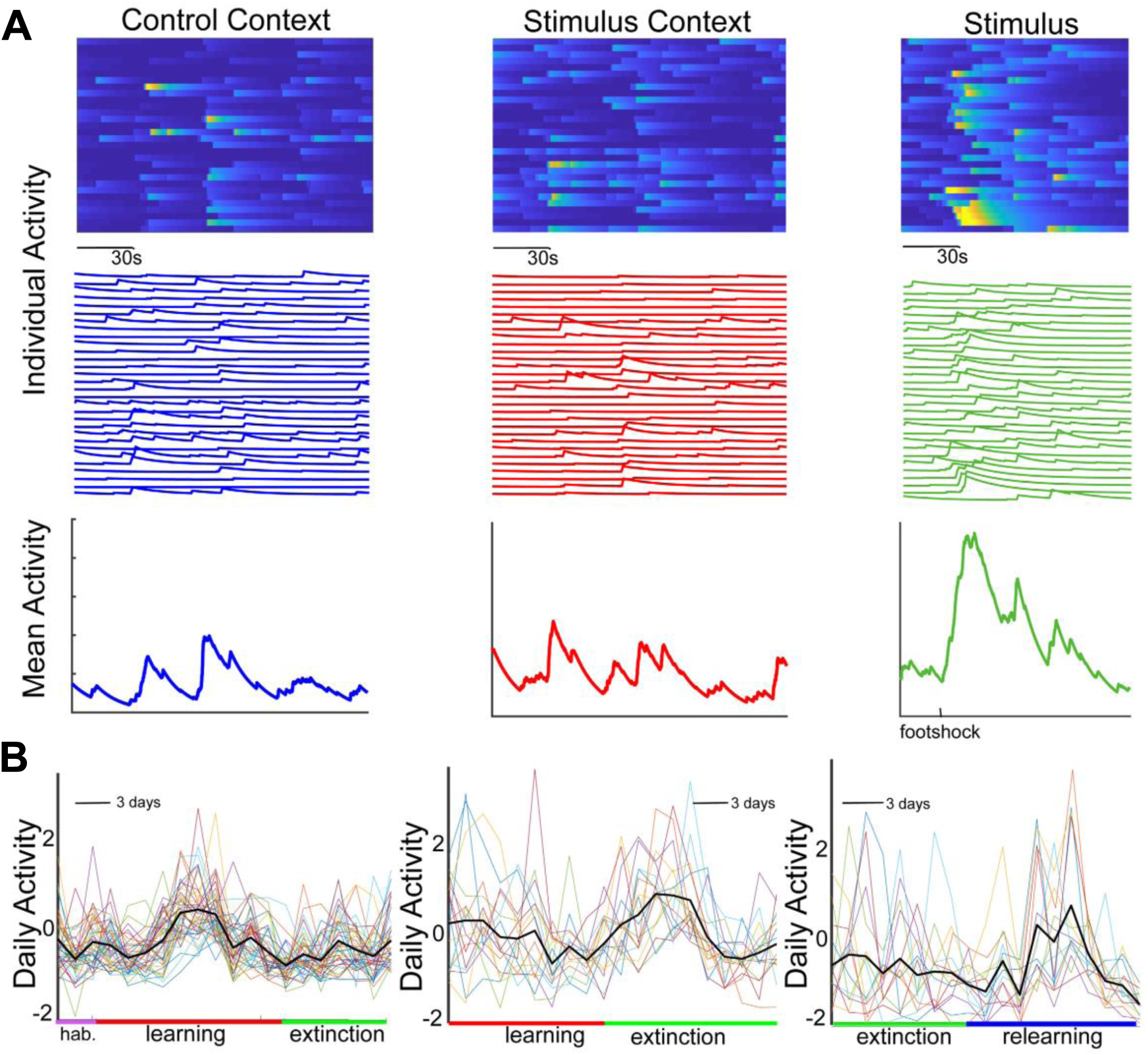
**A:** Neural activity from a single day during the learning phase of the experiment is split into three sections: control context (left column), stimulus context (before stimulus, middle column), and stimulus context (containing stimulus, right column). Calcium signal activity of individual neurons indicates little correlation between neural activity in the control and stimulus contexts. Neural activity shows a marked increase after application of the foot-shock stimulus, though this did not occur in all mice. **B:** Raw (unsmoothed) daily activity averages show experimental stage-related activity, though the effect is less differentiated before smoothing. (The activity shown here is the raw activity corresponding to Fig. 4F **(top)**).

## Supplemental Videos

**Video 1**: A simulated recording (left) is extracted by SCOUT into a product of spatial footprints and temporal traces (middle), forming a reconstruction of the recording with noise removed. The colored footprints in the extracted video (middle) have intensity given by their corresponding traces (right).

**Video 2**: SCOUT overcomes the difficult issues of non-rigid changes in spatial footprint size and location for consistent identification of neurons across sessions. Example tracked neurons are represented by colored ovals through 7 recording sessions.

